# p38γ and p38δ modulate innate immune response by regulating MEF2D activation

**DOI:** 10.1101/2022.12.09.519777

**Authors:** Alejandra Escós, Ester Díaz-Mora, Michael Pattison, Pilar Fajardo, Diego González-Romero, Ana Risco, José Martín-Gómez, Éric Bonneil, Nahum Sonenberg, Seyed Mehdi Jafarnejad, Juan José Sanz-Ezquerro, Steven C. Ley, Ana Cuenda

## Abstract

Evidence implicating p38γ and p38δ (p38γ/p38δ) in inflammation are mainly based on experiments using p38γ/p38δ deficient (p38γ/δ^-/-^) mice, which show low levels of TPL2, the kinase upstream of MKK1-ERK1/2 in myeloid cells. This could obscure p38γ/p38δ roles, since TPL2 is essential for regulating inflammation. Here we generated a p38γ^D171A/D171A^/p38δ^-/-^ (p38γ/δKIKO) mouse, expressing kinase-inactive p38γ and lacking p38δ. This mouse exhibited normal TPL2 levels, making it an excellent tool to elucidate specific p38γ/p38δ functions. p38γ/δKIKO mice showed a reduced inflammatory response and less susceptibility to LPS-induced septic shock and *Candida albicans* infection than wild-type mice. Gene expression analyses in LPS-activated WT and p38γ/δKIKO macrophages revealed that p38γ/p38δ regulated numerous genes implicated in innate immune response. Additionally, phospho-proteomic analyses and *in vitro* kinase assays showed that the transcription factor myocyte enhancer factor-2D (MEF2D) was phosphorylated at Ser444 via p38γ/p38δ. Mutation of MEF2D Ser444 to the non-phosphorylatable residue Ala increased its transcriptional activity and the expression of *iNOS* and *IL-1β* mRNA. These results suggest that p38γ/p38δ govern innate immune responses by regulating MEF2D phosphorylation and transcriptional activity.

## Introduction

The p38 mitogen-activated protein kinases (p38MAPK) are essential in cell adaptation to environmental changes, as well as in inflammatory cellular responses to pathogen infection (Cuenda & Rousseau, 2007; Cuenda & Sanz-Ezquerro, 2017). The stimulation of pattern-recognition receptors (PRRs), by pathogens or molecules from damaged cells, triggers the activation of p38MAPK and other signalling pathways that are critical to promote an innate immune response and production of inflammatory molecules (cytokines and other mediators) (Gaestel *et al*., 2009; Arthur & Ley, 2013). It is well established that among the four p38MAPK isoforms (p38α (*MAPK14*), p38β (*MAPK11*), p38γ (*MAPK12*) and p38δ (*MAPK13*)), p38α is an inflammation modulator and has been considered as a therapeutic target for inflammatory diseases and related pathologies (Gaestel et al., 2009; Arthur & Ley, 2013; Cuenda & Sanz-Ezquerro, 2017; Han *et al*., 2020). However, clinical trials using potent p38α inhibitors have failed (Cohen, 2009; Arthur & Ley, 2013).

p38γ and p38δ (p38γ/p38δ) are closely related kinases and have largely redundant functions, which makes it difficult to study the specific functions of these kinases using single knock-out mice. Recent studies have analysed mice lacking both p38γ and p38δ (p38γ/δ^-/-^) to demonstrate the importance of these kinases in the innate immune response and inflammation. p38γ/p38δ promote inflammation in several disease models, including sepsis, candidiasis, colitis, dermatitis, liver steatosis, arthritis, and cancer-associated with inflammation (Cuenda & Sanz-Ezquerro, 2017; Alsina-Beauchamp *et al*., 2018). Biochemical studies have indicated an important role for p38γ/p38δ MAPKs in myeloid cells (Risco *et al*., 2012; Cuenda & Sanz-Ezquerro, 2017; Alsina-Beauchamp *et al*., 2018). The production of cytokines and chemokines triggered by C-type lectin receptor (Dectin-1) and Toll like receptor (TLR) stimulation is reduced in macrophages lacking p38γ/p38δ (Alsina-Beauchamp *et al*., 2018). Also, compound p38γ/p38δ deficiency substantially reduces *TPL2* mRNA translation and TPL2 protein levels in macrophages (Risco *et al*., 2012; Alsina-Beauchamp *et al*., 2018; Escos *et al*., 2022). TPL2 is the key MAP3-kinase upstream of the MKK1-ERK1/2 in myeloid cells in innate immune responses. p38γ/δ^-/-^ macrophages show impaired TLR activation of Extracellular signal-Regulated Kinase 1/2 (ERK1/2) and reduced production of key TPL2-regulated cytokines such as TNFα (Risco *et al*., 2012; Alsina-Beauchamp *et al*., 2018), raising the possibility that the observed effects in p38γ/δ^-/-^ mice could be due in part to the reduced steady-state levels of TPL2. However, p38γ/p38δ deficiency does not completely recapitulate the effects observed in TPL2-deficient (TPL2^-/-^) mice suggesting specific p38γ/p38δ functions. For example, in a candidiasis mouse model, *Candida albicans* (*C. albicans*) infection and cytokine production is reduced in p38γ/δ^-/-^ mice, but increased in TPL2^-/-^ mice relative to wild-type (WT) (Alsina-Beauchamp *et al*., 2018). Also, in a chemically induced colitis-associated colon cancer model, deletion of p38γ/p38δ decreases tumour development, whereas in TPL2^-/-^ mice tumour development is increased (Koliaraki *et al*., 2012; Del Reino *et al*., 2014; Alsina-Beauchamp *et al*., 2018).

To investigate the specific TPL-2-independent functions of p38γ/p38δ in the innate immune responses, we generate a p38γ^D171A/D171A^/p38δ^-/-^ (p38γ/δKIKO) mouse that expressed a catalytically inactive p38γ isoform and lacks p38δ. Here, we report that, in contrast to p38γ/δ^-/-^ mice, p38γ/δKIKO mice show normal TPL2 protein levels, and therefore are a good tool to elucidate TPL2-independent roles of p38γ and p38δ *in vivo*. We studied p38γ/δKIKO mice inflammation and infection in two different mouse models of sepsis, and compared with WT mice. Also, we analysed the gene expression and protein phosphorylation in LPS-activated p38γ/δKIKO macrophages, and found that these were different compared to WT macrophages. We demonstrate that p38γ/p38δ phosphorylate the transcription factor myocyte enhancer factor-2D (MEF2D) at Ser444 (Ser437 in mouse), and that this phosphorylation modulates MEF2D’s transcriptional activity. MEF2D belongs to MEF2 family, is highly regulated by extracellular stimuli, and has been implicated in transcriptional control of cytokines in innate immune cells (Lu *et al*., 2021; Wang *et al*., 2021). Based on this result and other findings in this study, we demonstrate the importance of p38γ and p38δ for innate immune responses and suggest that pharmacological inhibition of p38γ and p38δ activity would be beneficial in inflammatory diseases and infection.

## Results and Discussion

### Generation and characterization of p38γ^D171A/D171A^, p38δ^-/-^ (p38γ/δKIKO) mice

Since p38γ expression in p38γ/δ^-/-^ cells restores TPL2 levels independently of its kinase activity (Risco *et al*., 2012; Escos *et al*., 2022), we decided to generate a p38γ/δKIKO mouse line by crossing p38γ^D171A/D171A^ and p38δ^-/-^ mice. p38γ/δKIKO mice will not have either p38γ or p38δ kinase activity as p38γ/δ^-/-^ mice. Genotype was confirmed by PCR (**Fig EV1A**). These mice, which have catalytically inactive p38γ and lack p38δ, were viable and fertile and had no obvious health problems. Western blot analysis using p38γ and p38δ antibodies confirmed that p38γ/δKIKO mouse embryonic fibroblasts (MEF) did not express p38δ and expressed p38γ, although to lower levels to that of WT cells (**Fig EV1B**). The expression levels of p38α, c-Jun N-terminal Kinase (JNK1/2) and extracellular signal-regulated kinase 5 (ERK5) were similar in p38γ/δKIKO and in wild type (WT) MEFs (**Fig EV1B**). In response to osmotic stress induced by sorbitol all signalling pathways were activated in p38γ/δKIKO and WT cells (**Fig EV1B**). Osmotic shock caused p38γ phosphorylation in both WT and p38γ/δKIKO MEF (**Fig EV1C**); however, phosphorylation of the protein hDlg at Ser158, a physiological p38γ substrate (Sabio *et al*., 2005) (*17*), was only observed in WT, but not in p38γ/δKIKO MEF (**Fig EV1D**), confirming that the p38γ expressed in p38γ/δKIKO cells is catalytically inactive.

We next examined Toll Like Receptor 4 (TLR4) stimulation by LPS in bone marrow derived macrophages (BMDM). Differentiation of bone marrow progenitor cells to macrophages was not affected in p38γ/δKIKO mice, as indicated by the expression of macrophage-specific membrane protein marker F4/80 (**Fig. EV2A**). We found that WT and p38γ/δKIKO BMDM exhibited similar levels of TPL2, whereas TPL2 expression was severely reduced in p38γ/δ^-/-^ cells as found previously (**Fig. 1A, 1B, Fig EV1E**). A20 Binding Inhibitor of NF-κB2 (ABIN2), a TPL2-associated protein, was expressed in WT and p38γ/δKIKO macrophages, but was not in p38γ/δ^-/-^ cells (**Fig. 1A**). As TPL2 mediates TLR activation of ERK1/2 signalling in macrophages (Gantke *et al*., 2011), we next analysed the activation of ERK1/2, as well as JNK1/2, p38α and the canonical NF-κB pathway, whose activation is triggered by TLR4 stimulation in macrophages (Lee & Kim, 2007; Gaestel *et al*., 2009; Takeuchi & Akira, 2010). ERK1/2 activation in p38γ/δKIKO BMDM was similar to that in WT cells (**Fig. 1A, 1B**). We also confirmed that the activation of ERK1/2 was impaired in p38γ/δ^-/-^ macrophages compared to WT (**Fig. 1A, 1B**). Phosphorylation of NF-κB1 p105 (p105) and activation loop phosphorylation of JNK1/2 and p38α were similar in all three LPS-stimulated WT, p38γ/δKIKO and p38γ/δ^-/-^ BMDM (**Fig. 1A**). These results indicate that TPL2 protein levels, and therefore activation of the ERK1/2 pathway by TLR4, were regulated by p38γ independently of its kinase activity. Accordingly, we have recently demonstrated that p38γ, and also p38δ, kinase-independent activity regulates the amount of TPL2 protein at two posttransciptional levels: 1) increasing TPL2 protein stability by binding to the complex TPL2/ABIN2/Nuclear Factor-κB1p105 (NF-κB1p105), and 2) modulating *TPL2* mRNA translation (Escos *et al*., 2022). The RNA binding protein aconitase-1 (ACO1) interacts with the 3’ untranslated region (UTR) of *TPL2* mRNA, repressing its translation and decreasing the levels of TPL2 protein in the cells (Escos *et al*., 2022). In the absence of p38δ, p38γ interacts with ACO1 impairing its association with *TPL2* 3’UTR, which leads to an increase in *TPL2* mRNA translation (Escos *et al*., 2022).

**Figure 1.**
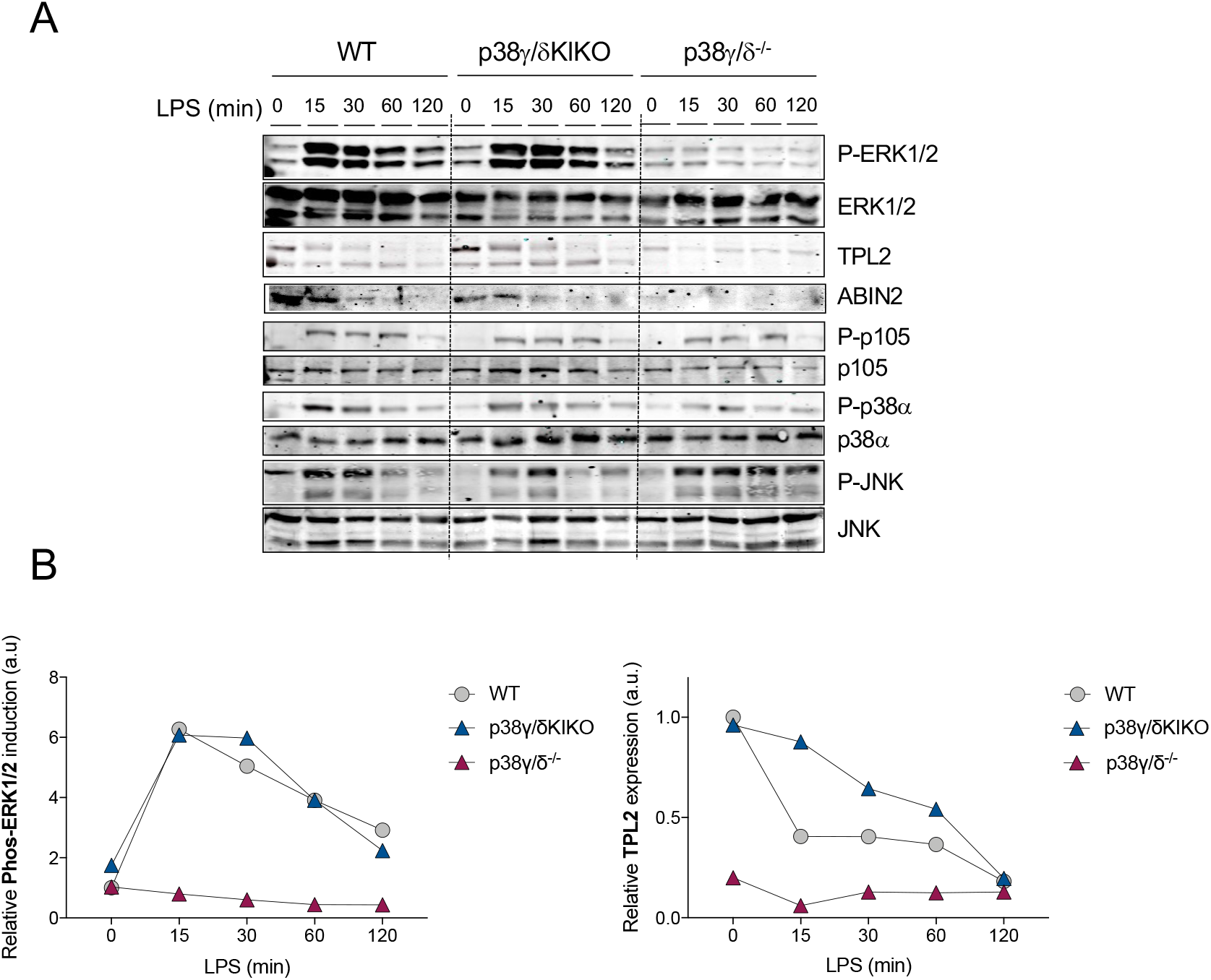
LPS-induced ERK1/2 activation in p38γ/δKIKO macrophages. **(A)** BMDM from WT, p38γ/δ^-/-^, and p38γ/δKIKO mice were exposed to LPS (100 ng/ml) for the indicated times. Cell lysates (30 μg) were immunoblotted with the indicated antibodies to active phosphorylated p38α (P-p38α), JNK1/2 (P-JNK) and ERK1/2 (P-ERK1/2). Phosphorylated p105 (P-p105) and total protein levels of p38α, JNK1/2 (JNK), ERK1/2, TPL2 and p105 were also measured in the same lysates. Representative blots of two independent experiments with similar results are shown. **(B)** Intensity of the bands corresponding to Phos-ERK1/2 and TPL2 (panel A) were quantified. Relative data to WT Time 0 min are shown.

### p38γ/δKIKO mice are protected from *Candida albicans* infection and LPS-induced septic shock

We next used p38γ/δKIKO mice to determine the role of p38γ and p38δ signalling in innate immune responses and inflammation. We performed a comparative analysis of *C. albicans* infection in p38γ/δKIKO and WT mice. Fungal burden and total number of infiltrating leucocytes (CD45^+^ cells) in the kidneys of p38γ/δKIKO mice were lower than in WT control at day 3 post-infection (**Fig. 2A, 2B**). We then checked that resting levels of both macrophages and neutrophils were similar in the bone marrow (BM) and spleen of WT and p38γ/δKIKO mice (**Fig. EV2B-D)**. Absolute cell numbers in BM and spleen, as well as spleen weight, were similar in both genotypes (**Fig. EV2E, EV2F**). Next, we determined the renal recruitment of macrophages (F4/80^+^ cells) and neutrophils (Ly6G^+^ cells), the major leukocyte types involved in *C. albicans*-induced inflammation (Alsina-Beauchamp *et al*., 2018). We found that after *C. albicans* infection, the percentage and the total number of F4/80^+^ cells were significantly reduced in p38γ/δKIKO mice compared to WT, whereas Ly6G^+^ cells recruitment was also reduced but not significantly different between genotypes (**Fig. 2C**).

**Figure 2.**
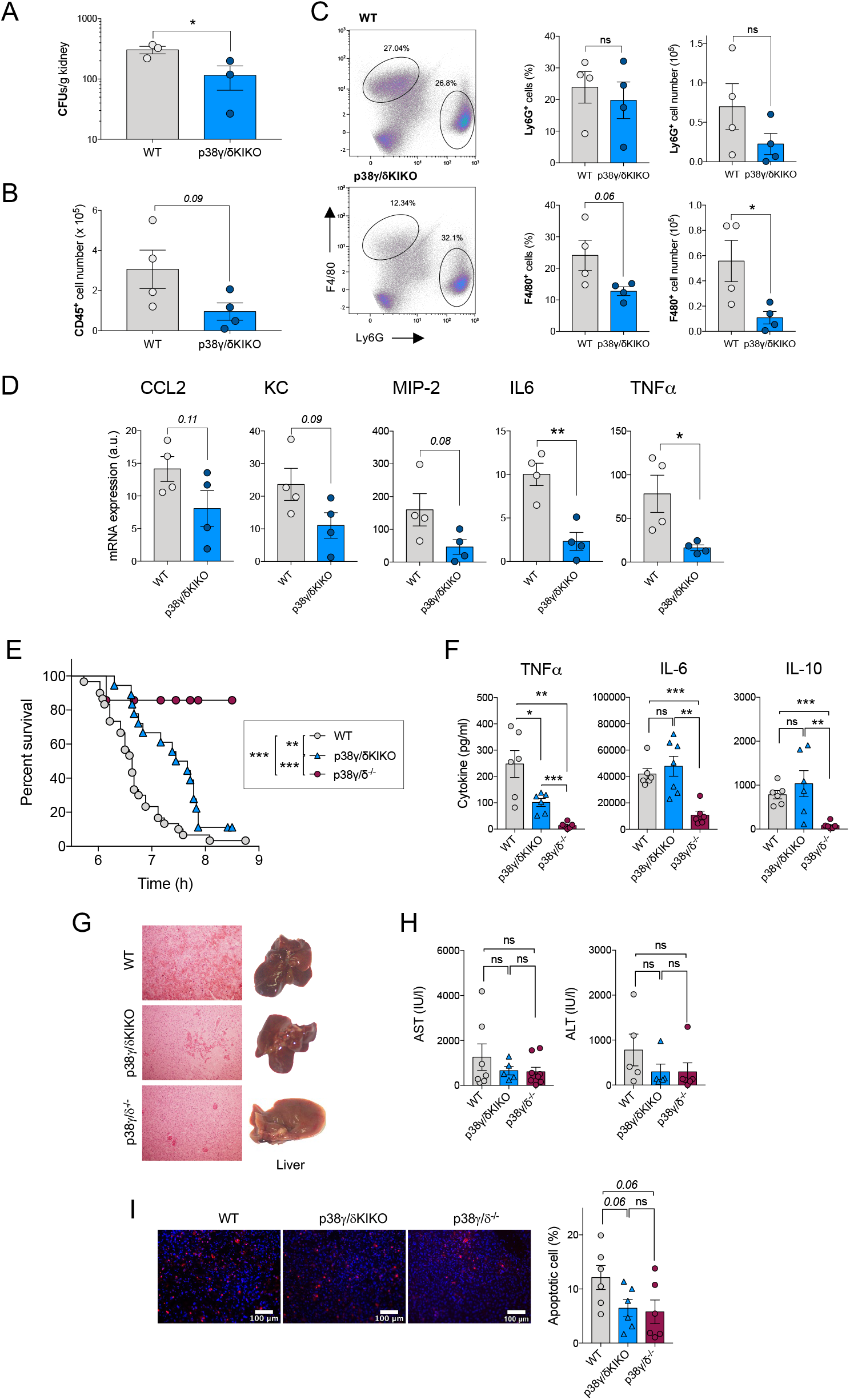
Reduced inflammation in p38γ/δ KIKO mice in response to septic shock. (**A**) WT and p38γ/δKIKO mice were intravenously injected with 1 × 10^5^ CFU of *C. albicans*. Kidney fungal load was determined 3 days after infection. Each symbol represents an individual mouse. Figure shows mean ± SEM, ns not significant; \**p* ≤0.05, relative to WT kidney cells. (**B, C**) Kidney cells were stained with (**B**) anti-CD45, (**C**) anti -Ly6G and -F4/80 antibodies and positive cells analysed by flow cytometry. CD45+ cells were gated and -F4/80+ and -Ly6G+ cells analysed by flow cytometry. Representative profiles are shown. Each symbol represents an individual mouse. Histograms shows mean ± SEM, ns not significant; \**p* ≤0.05, relative to WT kidney cells. **(D)** Mice were treated as in (A) and the mRNA levels of indicated genes in the kidney were measured by qPCR 3 days after infection. Each symbol represents an individual mouse. Figure shows mean ± SEM (*n* = 4 mice/condition). ns, not significant, \**p* ≤0.05, \*\**p* ≤0.01. **(E)** WT (*n* = 31), p38γ/δ^-/-^ (*n*= 18) and p38γ/δKIKO (*n* = 19) mice were injected with LPS (50 μg/kg) and D-Gal (1g/Kg), and survival was monitored for up to 9 hours. Graph shows % survival at the indicated times. ** *p* ≤0.01, **\***\*\* *p* ≤0.001. **(F)** Serum from mice (E) was collected 2 h after LPS/D-Gal injection, and TNFα, IL-6 and IL10 were measured in a Luminex cytokine assay. Each symbol represents an individual mouse. Figure shows mean ± SEM (*n* = 6-7 mice). ns, not significant; \**p* ≤ 0.05, \*\**p* ≤ 0.01, \*\*\**p* ≤ 0.001. **(G)** Livers were collected 6 h after LPS/D-Gal injection. Panels show H&E stained liver sections (left) and whole livers (right). **(H)** Serum ALT (alanine transaminase) and AST (aspartate aminotransferase) activity at 6 h after LPS/D-Gal injection. Each symbol represents an individual mouse. Figure shows mean ± SEM (*n* = 5-7 mice). ns, not significant. (**I**) Apoptotic TUNEL positive (red) and total nuclei (Hoechst stained-blue) cells were counted using ImageJ programme and the percentage of apoptotic cells calculated. 25 sections per mouse were scored. Representative TUNEL stained liver sections are shown, and figure shows mean ± SEM (*n* = 6 mice). ns, not significant. Each symbol represents an individual mouse.

The decrease in renal recruitment of leukocytes was paralleled with a reduced chemokine and cytokine expression (**Fig 2D**) as found previously in p38γ/δ^-/-^ mice (Alsina-Beauchamp *et al*., 2018). Cytokines *IL-1β, TNFα* and *IL-6* mRNA levels were clearly lower in the kidney of p38γ/δKIKO mice than in WT animals at day 3 post-infection (**Fig 2D**). Together, these findings support the role of p38γ/p38δ in the inflammatory response to *C. albicans* by modulating the production of inflammatory molecules and the recruitment of leukocytes into the *C. albicans* infected kidney. These data also show that p38γ/δKIKO behave similar to p38γ/δ^-/-^ mice in a candidiasis model, as described in (Alsina-Beauchamp *et al*., 2018), and validate p38γ/δKIKO mice as a new tool to study the physio-pathological function of alternative p38MAPK *in vivo*, independently of TPL2.

Since TPL2 does not play a critical role in *C. albicans* infection (Alsina-Beauchamp *et al*., 2018), we also examined if p38γ/δKIKO and p38γ/δ^-/-^ mice behave similarly in a TPL2-dependent septic shock model induced by the endotoxin lipopolysaccharide (LPS) (Dumitru *et al*., 2000). We have previously reported that p38γ/p38δ deficiency protects mice from sepsis induced by LPS (Risco *et al*., 2012). We then treated mice with *Escherichia coli*-derived LPS plus the hepatotoxic compound D-Galactosamine (D-Gal) and checked mouse survival comparing p38γ/δKIKO with WT or p38γ/δ^-/-^ mice. p38γ/δKIKO mice were significantly more resistant to LPS-induced septic shock than WT mice, although the compound p38γ/p38δ deficiency had a more pronounced protective effect (**Fig. 2E**). We have already shown that the reduced susceptibility of p38γ/δ^-/-^ mice to LPS-induced septic shock was at least in part due to an overall decrease in cytokine production (Risco *et al*., 2012). We observed significantly reduced levels of circulating cytokines IL-6, TNFα and IL-10 in p38γ/δ^-/-^ mice than in WT mice after LPS/D-Gal treatment, whereas in p38γ/δKIKO mice only the production of TNFα was significantly reduced compared to WT (**Fig. 2F**). Haemorrhage in liver from p38γ/δKIKO mice was lower than WT, but was higher than that observed in p38γ/δ^-/-^ liver (**Fig. 2G**). The circulating serum transaminases, alanine transaminase (ALT), and aspartate aminotransferase (AST), two markers of hepatic necrosis, were noticeably higher in WT than in p38γ/δ^-/-^ and p38γ/δKIKO mice, although this difference was not statistically significant due to mouse individual variation (**Fig. 2H**). In addition, TUNEL analysis showed a decreased apoptosis in p38γ/δ^-/-^ and p38γ/δKIKO mice compared to WT (**Fig. 2I**), indicating a reduction in liver cell death and protection against liver damage in the mutant mice.

These comparative experiments in p38γ/δKIKO and p38γ/δ^-/-^ mice show that the combined deletion of p38γ/p38δ has a more pronounced impact in LPS-induced cytokine production in mice. This indicates that the effects observed in p38γ/δ^-/-^ mice are partly due to the decreased TPL2 levels and the blockade of ERK1/2 pathway signalling, and also confirms a role for p38γ/p38δ, independently of TPL2 expression, in regulating LPS/D-Gal-induced septic shock by increasing liver damage and TNFα production, and in *C. albicans* infection modulating the inflammatory response.

### Analysis of TLR4-induced gene expression in p38γ/δKIKO macrophages

Since p38γ/p38δ are important in the immune response to septic shock and to the TLR4 ligand LPS in the macrophage (Risco *et al*., 2012; Alsina-Beauchamp *et al*., 2018), we stimulated WT and p38γ/δKIKO BMDM with LPS and analysed gene expression by RNA-sequencing (RNA-Seq) to identify target genes specifically affected by the lack of p38γ kinase activity and p38δ expression. Both, WT and p38γ/δKIKO BMDM, showed a strong transcriptional response at 30 and 60 min of stimulation with LPS (**Fig. 3A-C**). WT showed increased (at least 2.0-fold, *p*-value <0.05) expression of 197 and 533 genes at 30 and 60 min, respectively, whereas p38γ/δKIKO BMDM showed increased expression of 264 and 647 genes at 30 and 60 min, respectively (**Fig. 3A**). Venn diagram analysis revealed that ∼60% of the up-regulated genes were common between WT and p38γ/δKIKO BMDM at both 30 and 60 min; whereas ∼30%-40% of the down-regulated genes were shared between genotypes (**Fig. 3C**).

**Figure 3.**
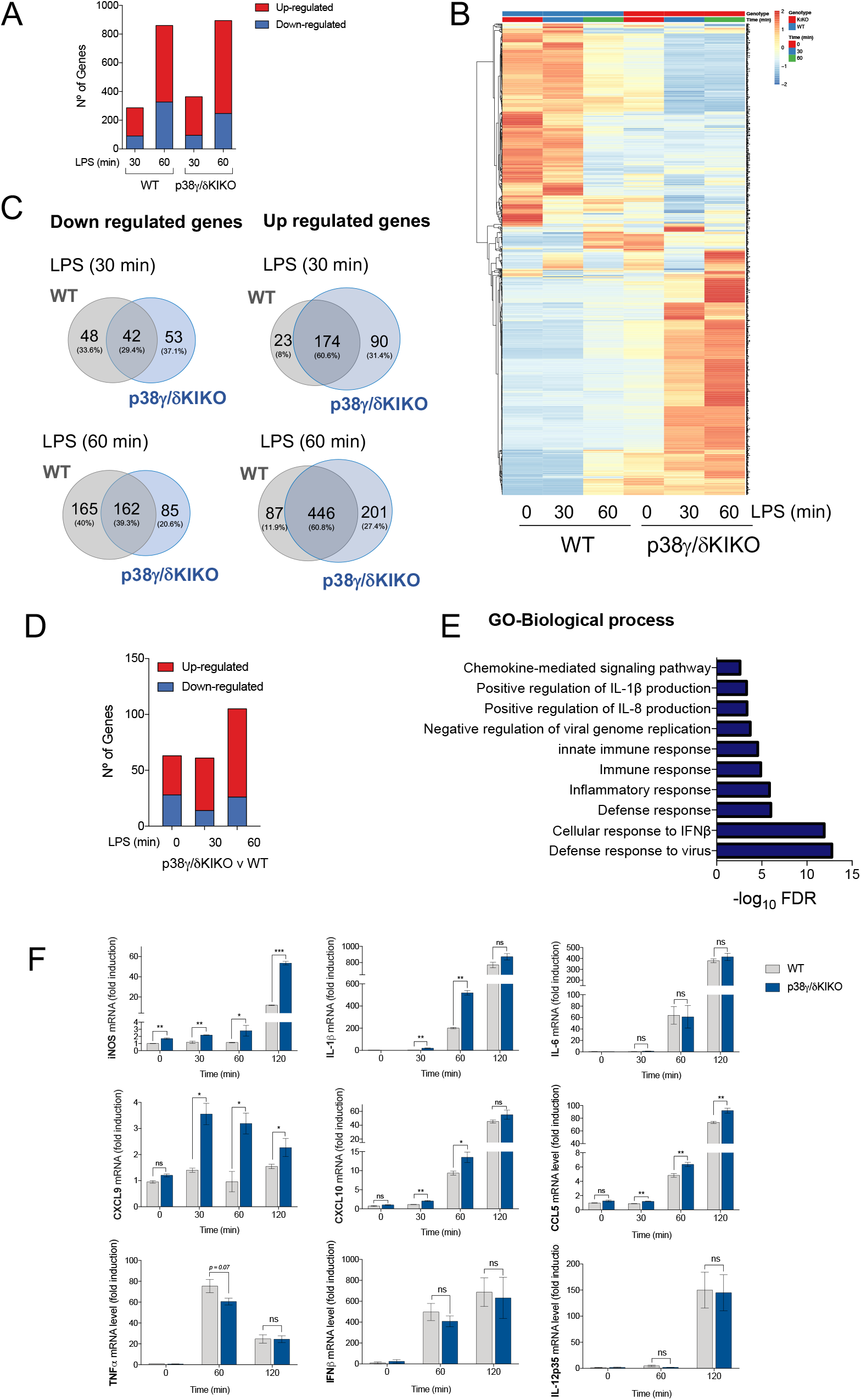
RNA-sequencing analysis in LPS-stimulated WT and p38γ/δKIKO macrophages. **(A)** BMDM from WT and p38γ/δKIKO mice were exposed to LPS (100 ng/ml) for 0, 30 and 60 min and gene expression analysed by RNA sequencing. Bar plot showing the number of differentially expressed genes up- or down-regulated after LPS stimulation (−2< logFC <2, *p*-value <0.05). Three samples per condition were used. **(B)** Hierarchical heatmap of the differentially expressed genes in panel (A). **(C)** Venn diagrams showing the overlaps of genes upregulated or downregulated over the time course of LPS stimulation in p38γ/δKIKO and WT macrophages. **(D)** Bar plot showing the number of differentially expressed genes up- or down-regulated (−1.5< logFC <1.5, *p*-value <0.05) in p38γ/δKIKO macrophages compared with WT after LPS stimulation at the indicated times (minutes). Three samples per condition were used. **(E)** Enrichment analysis of GO biological processes of the differentially expressed genes in LPS-activated p38γ/δKIKO macrophages after LPS stimulation. **(F)** BMDM from WT and p38γ/δKIKO mice were exposed to LPS (100 ng/ml) for the indicated times. mRNA expression of indicated genes at different times, relative to WT expression at 0 h, was determined by qPCR and normalised to β-actin mRNA. Data show mean ± SEM (*n* = 3-6). ns, not significant; \**p* ≤0.05; \*\**p* ≤0.01, \*\*\**p* ≤ 0.001, relative to WT mice in the same conditions.

We next compared gene expression profiles between genotypes in LPS-stimulated p38γ/δKIKO and WT BMDM (**Fig. 3D**) and selected genes with either increased or decreased expression between genotypes by at least 1.5-fold (−1.5< logFC <1.5), with a *p*-value <0.05 (**Fig. 3D, Table EV1**). We found that overall, in p38γ/δKIKO macrophages the expression of 54 genes was downregulated and the expression of 120 genes was upregulated as compared to WT (**Fig. EV3A)**. Different gene profiles of p38γ/δKIKO and WT BMDM at basal conditions indicates that p38γ and p38δ also regulate gene expression in resting conditions. Gene Ontology (GO) analyses of differentially expressed genes using DAVID (the Database of Annotation, Visualization and Integrated Discovery) revealed the enrichment for genes implicated in the innate immune and inflammatory response (**Fig. 3E, Fig. EV3B**).

One of the main processes affected in LPS-stimulated p38γ/δKIKO was the immune response and cytokine production. Therefore, to validate the RNA-Seq analysis we performed a comparative analysis by qPCR of the expression of genes involved in these processes (**Table EV1, Fig. EV3B, Fig. 3F**). We found that after LPS treatment the expression of *CCL5, CXCL9* and *iNOS* mRNAs increased in p38γ/δKIKO macrophages compared to WT, as observed in RNA-Seq analysis. *IL1β* and *CXCL10* mRNA expression was also significantly higher at 30 and 60 min after LPS treatment in p38γ/δKIKO BMDM. Whereas, *IFNβ* and *IL-12p35* mRNA expression, which are TPL2-ERK1/2 targets, and *IL-6* and *TNFα* mRNA expression were similar in p38γ/δKIKO and WT macrophages (**Fig. 3F**). These results indicate that p38γ and p38δ in macrophages specifically regulate production of inflammatory molecules in response to LPS.

We also assessed cytokines expression levels using a Mouse Cytokine Array, at 6 h after LPS treatment. We found that the global cytokine expression pattern in WT and p38γ/δKIKO macrophages seemed similar (**Fig. EV3C**); nonetheless, after quantification we found that the levels of CXCL1, CCL2, CCL12, CXCL2, CCL5, IL-5, IL-10, sICAM-1, IL-12p70 and IL-1α were significantly higher in p38γ/δKIKO cells (**Fig. EV3D**). We confirmed that IL-1β production was increased in p38γ/δKIKO macrophages compared to WT, whereas TIMP-1, TREM-1 and IL-6 were decreased (**Fig. EV3D**). These data indicate that p38γ and p38δ control the production of inflammatory molecules in response to LPS in macrophages by modulating cytokine transcription and synthesis.

### Identification of p38γ and p38δ-dependent protein phosphorylation

To establish the molecular mechanism by which p38γ/p38δ modulate macrophage activation and cytokine production, we sought to identify the p38γ/p38δ substrates by performing a comparative phosphoproteomic analysis of LPS-stimulated WT, p38γ/δKIKO, and p38γ/δ^-/-^ macrophages. We used peritoneal macrophages since p38γ and p38δ are expressed at much higher levels in these macrophages than in BMDM (Risco *et al*., 2012). We first confirmed with western blot that ERK1/2 phosphorylation was similar in WT and p38γ/δKIKO cells but was impaired in p38γ/δ^-/-^, and that p38α activation was similar in all LPS-stimulated macrophages (**Fig. EV4A**). Additionally, the LPS-induced cytokine production in peritoneal macrophages was comparable to that of BMDM (**Fig. EV4B**).

Comparison of the phosphoproteomes of LPS-stimulated WT, p38γ/δ^-/-^ and p38γ/δKIKO with control unstimulated macrophages confirmed the phosphorylation of proteins from the classical/canonical TLR4-activated signalling pathways such as p38α, its substrate the kinase MAPKAPK2 (MK2) and the protein Tristetraproline (TTP), which is itself a MK2 substrate (**Fig. EV4C**). This indicates the robustness of our experimental approach. Heatmap analysis revealed that peptide phosphorylation patterns were similar in p38γ/δ^-/-^ and p38γ/δKIKO LPS-stimulated macrophages, as compared to WT (**Fig. 4A**). Comparison with the WT showed significant reduction of 35 and 16 phosphopeptides in p38γ/δKIKO and p38γ/δ^-/-^ LPS-treated samples, respectively (−1.0< logFC <1.0, with a *p*-value <0.05) (**Fig. 4B-D**). Notably, we observed that 158 and 94 phosphopeptides were increased in p38γ/δKIKO and p38γ/δ^-/-^ samples, respectively (−1.0< logFC <1.0, with a *p*-value <0.05) (**Fig. 4B-D**). The number of upregulated phosphopeptides was larger than those downregulated indicating that, upon LPS-stimulation, p38γ and p38δ control the activity of a yet unidentified kinase, or phosphatase, or both, in macrophages. Venn diagram analysis revealed that only the phosphorylation of peptides from nucleolar protein 56 (Nop56), Myocyte enhancer factor 2D (MEF2D) and osteopontin (spp1), was downregulated in both p38γ/δKIKO and p38γ/δ^-/-^ macrophages (**Fig. 4D**).

**Figure 4.**
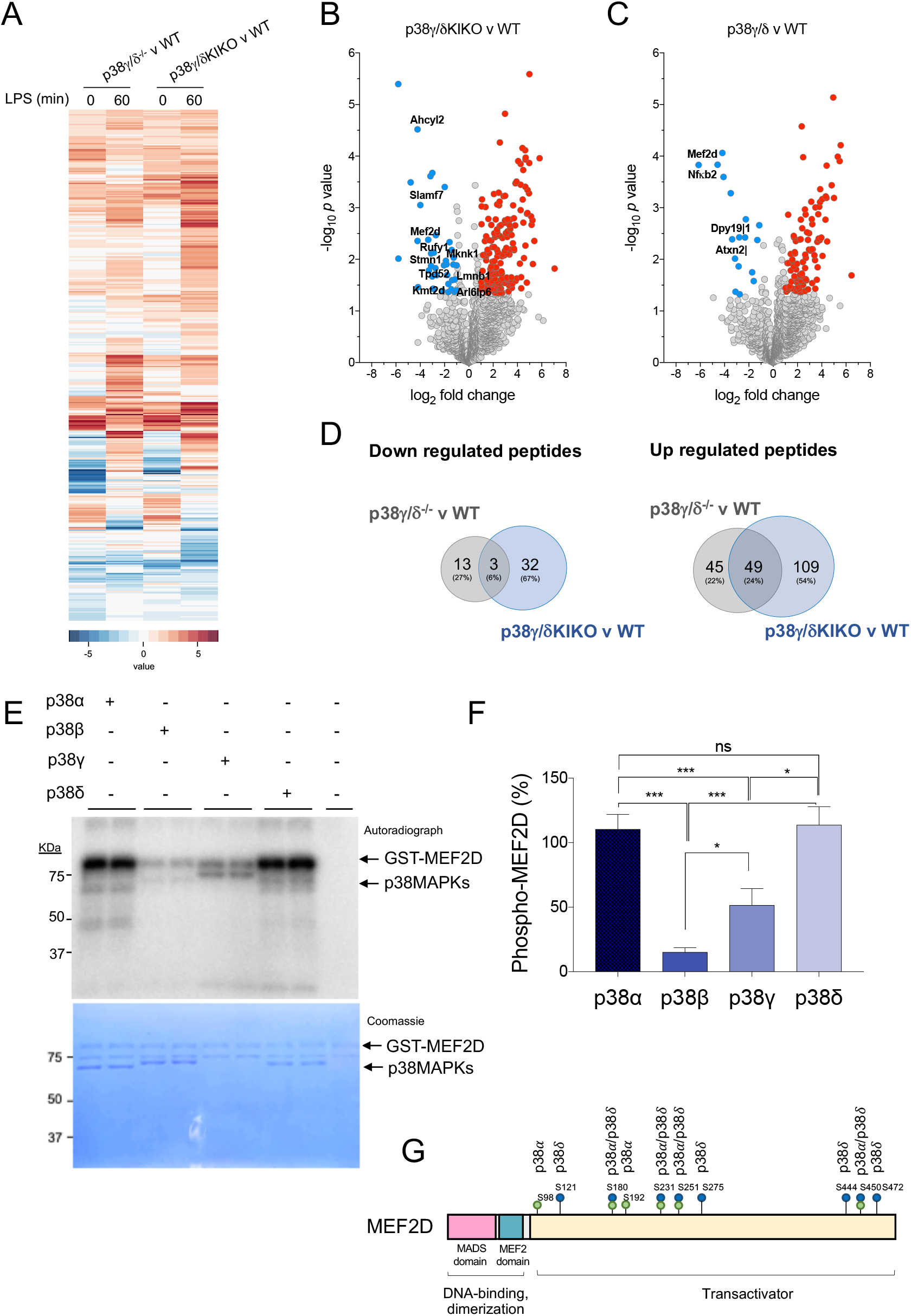
Identification of proteins phosphorylated by p38γ/p38δ. (**A**) Peritoneal macrophages from WT, p38γ/δ^-/-^ and p38γ/δKIKO mice were exposed to LPS (100 ng/ml) for 0 or 60 min and phosphorylated peptides identified. Heatmap showing differentially phosphorylated phosphosites up- or down-regulated after LPS stimulation. (**B, C**) Volcano plots showing the differential phosphoproteome and statistical significance between (**B**) p38γ/δKIKO and WT (*n* = 3), and (**C**) p38γ/δ^-/-^ and WT (*n*= 3) macrophages treated with LPS for 60 min. Examples of significantly enriched up-regulated (red) and down-regulated (blue) phosphosites are shown (−1.0< logFC <1.0, *p*-value <0.05). **(D)** Venn diagrams showing the overlaps of differentially phosphorylated proteins (upregulated or downregulated) at 60 min LPS stimulation in p38γ/δKIKO and p38γ/δ^-/-^ macrophages compared to WT. **(E)** Recombinant GST-MEF2D (or Myelin basic protein (MBP) as control) was incubated with active recombinant p38α, p38β, p38γ or p38δ for 60 min in a phosphorylation reaction mix containing Mg-[γ^32^P]ATP, as described in the materials and methods. The activity of recombinant p38α, p38β, p38γ or p38δ was matched using MBP as substrate and 0.5 U/ml were used in the assay. Reaction was stopped with SDS-sample buffer. Samples were resolved in SDS-PAGE, subjected to Coomassie blue staining and autoradiography. **(F)** Bands corresponding to ^32^P-MEF2D from panel (E) were quantified. Data show mean ± SEM from two experiments in duplicate. ns, not significant \**p* ≤0.05; \*\*\**p* ≤ 0.001, relative to MEF2D phosphorylation by p38α. **(G)** Schematic representation of the sites in MEF2D phosphorylated by p38α and/or p38δ, all of them located in the transactivation domain.

We next examined the downregulated phosphorylation sites lying within the well-characterized p38MAPK phosphorylation site consensus S/T-P motif (Cuenda & Rousseau, 2007), as these represent likely candidates for direct p38γ and p38δ physiological substrates. We identified 4 candidates with S/T-P phosphorylation site in (p38γ/δ^-/-^ v WT) comparison, and 11 in (p38γ/δKIKO v WT) comparison, one of them being the previously described p38δ substrate, Stathmin (Stmn1) (Cuenda & Rousseau, 2007) (**Table EV2**). MEF2D (Ser437 in mouse and Ser444 in human) was the only protein identified in both (p38γ/δ^-/-^ v WT) and (p38γ/δKIKO v WT) comparisons. MEF2D is a member of the transcription factor MEF2 family, which contains MEF2A, B, C and D and were initially identified in muscle, but are also expressed in neurons, and cells of the immune system such as T and B cells and myeloid cells (McKinsey *et al*., 2002).

The activity of the MEF2 transcription factors is regulated by p38α-mediated phosphorylation (McKinsey *et al*., 2002); however, to our knowledge, the phosphorylation of MEF2D by p38γ and/or p38δ in cells has not been demonstrated. We then checked if MEF2D was phosphorylated by p38γ and/or p38δ using γ^32^P-ATP in *in vitro* kinase assay, and compared with the phosphorylation by p38α and p38β. We found that MEF2D was equally phosphorylated by p38α and p38δ, and less by p38γ, whereas p38β weakly phosphorylated MEF2D (**Fig. 4E-F**). We determined the MEF2D sites phosphorylated by p38α and p38δ, *in vitro*, by mass spectrometry analysis (**Fig. EV5A, B**). We identified ten amino acid phosphorylated by these p38MAPKs, although the p38δ phosphorylation sites within MEF2D differed from those phosphorylated by p38α (**Table EV3**). Four of the sites were phosphorylated by the two kinases, whereas Ser98 and Ser192 were exclusively phosphorylated by p38α, and Ser121, Ser275, Ser444 and Ser472 were specific for p38δ (**Table EV3, Fig. 4G, Fig EV5B**). This confirmed that Ser444 (Ser 437 in mouse) was one of MEF2D phosphorylation sites specific for p38δ.

### p38γ and p38δ regulate MEF2D transcriptional activity

MEF2D is activated by LPS, and regulates inflammation by modulating the expression of proinflammatory factors. Thus, MEF2D decreases NLRP3, iNOS and IL-1β expression in microglia, and IL-10 transcription in macrophages (Yang *et al*., 2015; Pattison et al., 2020; Lu *et al*., 2021). Transcriptional activity of MEF2D can be partially modulated by post-translational modifications such as phosphorylation. However, which kinases regulate MEF2D phosphorylation and function are still largely unknown. To investigate whether p38γ and p38δ modulate MEF2D transcriptional activity, we examined the TLR4-induced expression of *c-Jun, CD14*, and *HDAC7* genes, whose transcription is regulated by MEF2D (Han *et al*., 1995; Park *et al*., 2002; Lu *et al*., 2021). qPCR analysis showed that LPS-induced *c-Jun, CD14* and *HDAC7* mRNAs was significantly increased in p38γ/δKIKO macrophages compared with WT cells (**Fig. 5A**).

**Figure 5.**
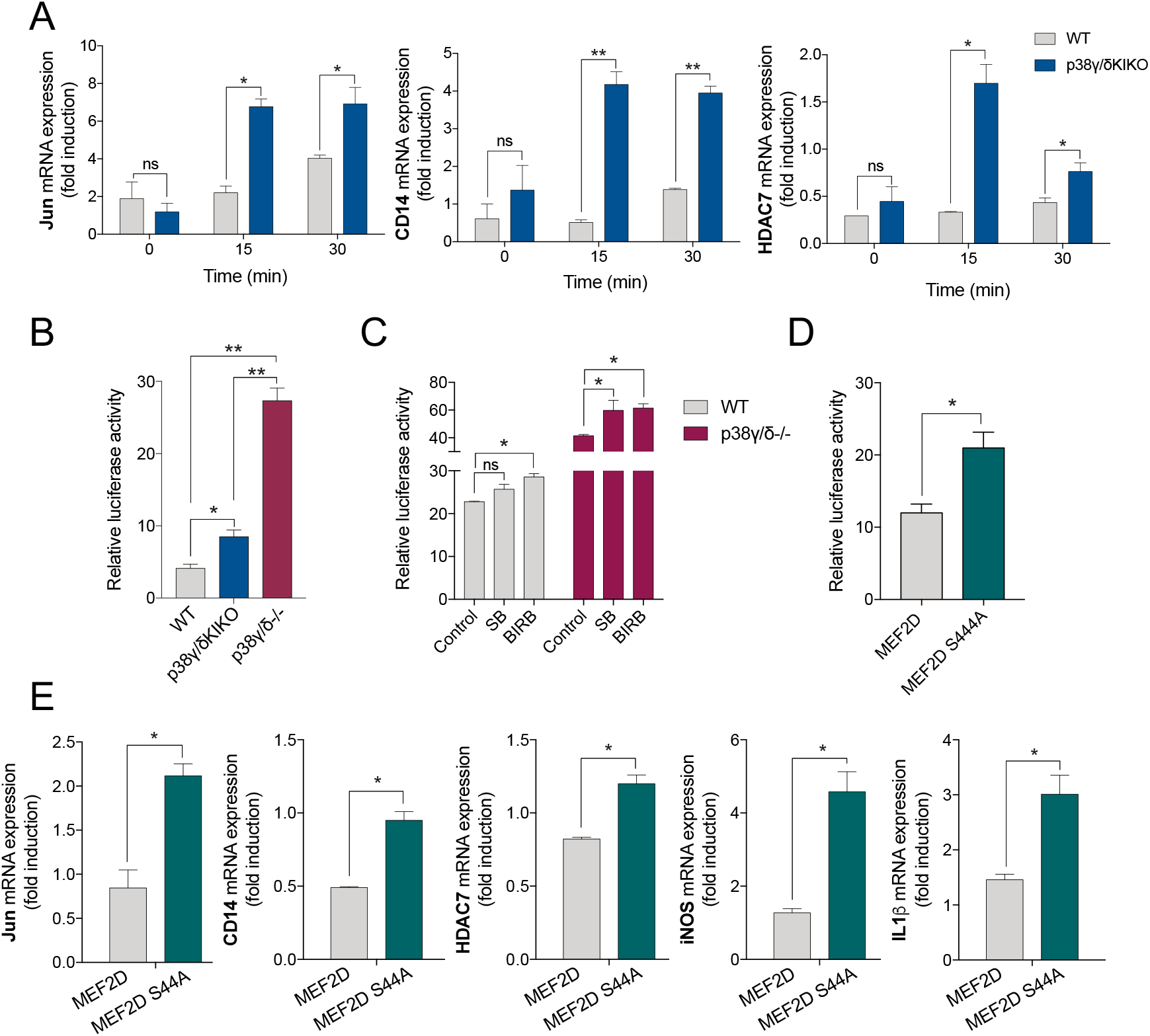
Regulation of MEF2D transcriptional activity by p38γ and p38δ and S444 phosphorylation. **(A)** BMDM from WT and p38γ/δKIKO mice were exposed to LPS (100 ng/ml) for the indicated times. Relative mRNA expression of *Jun, CD14* and *HDAC7* genes at different times was determined by qPCR. Results were normalized to *β-actin* RNA expression and fold induction was calculated relative to WT expression at 0 min. Figure shows mean ± SEM from two experiments in triplicate. ns, not significant; \**p* ≤0.05 and \*\**p* ≤0.01, relative to WT mice in the same conditions. **(B)** WT, p38γ/δKIKO and p38γ/δ^-/-^ fibroblasts were co-transfected with three plasmids coding for: Flag-MEF2D, Renilla and luciferase firefly gene under the control of MEF2 response elements. Luciferase activity values were normalised against Renilla. Figure shows mean ± SEM from one representative experiment in triplicate. This experiment was repeated twice with similar result. ns, not significant; \**p* ≤0.05; \*\**p* ≤0.01 and \*\*\**p* ≤ 0.001. **(C)** WT and p38γ/δ^-/-^ fibroblasts were transfected as in (B). Cells were incubated with the indicated p38MAPK inhibitor (or DMSO as control) for 6 h before lysis. Luciferase activity was calculated as in (B). **(D)** WT fibroblasts were transfected with plasmid Flag-MEF2D or Flag-MEF2D^S444A^, plus plasmids coding Renilla and luciferase firefly gene under the control of MEF2 response elements. Luciferase activity values were normalised against Renilla. Figure shows mean ± SEM from one representative experiment in triplicate. \**p* ≤0.05. **(E)** WT fibroblasts were transfected with Flag-MEF2D or Flag-MEF2D^S444A^, and stimulated with LPS for 30 min. Relative mRNA expression of the indicated genes was determined by qPCR. Results were normalized to *β-actin* RNA expression and fold induction was calculated relative to MEF2D expression. Figure shows mean ± SEM from two experiments in triplicate. \**p* ≤0.05.

To further examine the possible role of p38γ/p38δ in the specific transcriptional activity of MEF2D we performed reporter assays using a plasmid encoding a *Firefly Luciferase* gene under the control of MEF2 response elements. p38γ/δKIKO, p38γ/δ^-/-^ and WT fibroblasts were transfected with the reporter plasmid and a plasmid encoding Flag-MEF2D. We found a marked increase in luciferase activity in p38γ/δ^-/-^ and p38γ/δKIKO cells compared to WT cells (**Fig. 5B**). Interestingly, we observed that the transcriptional activity of MEF2D was significantly much higher in p38γ/δ^-/-^ than in p38γ/δKIKO cells suggesting that, in addition to phosphorylation, the interaction of p38γ with MEF2D or with other protein might be affecting MEF2D transcriptional activity; this question could be addressed in future works. Next, we analysed the effect of the p38MAPK inhibitor BIRB796 on MEF2D transcriptional activity. Since this compound inhibits all four (p38α, p38β, p38γ and p38δ) p38MAPK isoforms, we also used in parallel the compound SB203580 that blocks p38α and p38β only (Kuma *et al*., 2005), so we could determine which effect was caused by p38α/p38β or by p38γ/p38δ inhibition. Incubation with BIRB796, but not with SB203580 significantly increased luciferase activity in WT cells (**Fig. 5C**), indicating that p38γ and p38δ activity regulates MEF2D-mediated transcription. In p38γ/δ^-/-^ cells, luciferase activity was significantly and equally increased by SB20358 and BIRB796, suggesting that p38α can control MEF2D activity in the absence of p38γ/p38δ.

MEF2D transcriptional activity is modulated by phosphorylation, and phosphorylation at different MEF2D sites has different effects in its transcriptional activity (Tang *et al*., 2005; Ke *et al*., 2015). For example, MEF2D transcriptional activity is increased after phosphorylation at Thr259, Ser275, Ser294 and Ser314 by the kinase Ataxia telangiectasia mutated (ATM) in neurons (Chan *et al*., 2014), or at Ser179 by ERK5 in Hela cells (Kato *et al*., 2000). However, phosphorylation of MEF2D at Ser251 by DYRK1, in HEK293 cells (Wang *et al*., 2021), or of Ser444 by Cdk5, has been implicated in the repression of MEF2D transcriptional activity in neurons (Tang *et al*., 2005). MEF2D-Ser444 was phosphorylated *in vitro* by p38δ, and phosphorylation at this residue was decreased in p38γ/δ^-/-^ and p38γ/δKIKO cells compared to WT cell (**Fig. 4B-C**). Consistently, the expression of *c-Jun, CD14, HDAC7, iNOS* and *IL-1β* mRNA, which are MEF2D-regulated genes (Han *et al*., 1995; Park *et al*., 2002; Lu *et al*., 2021), is increased by the lack of p38γ/p38δ activity. We then mutated Ser444 to Ala to generate MEF2D-Ser444Ala (MEF2D^S444A^) mutant and study its effect on MEF2D transcriptional activity in WT cells. Transfection of MEF2D^S444A^ mutant caused a significant increase in luciferase activity compared to MEF2D (**Fig. 5D**), similar to that observed in p38γ/δKIKO cells (**Fig. 5B**). In addition, qPCR analysis showed that *c-Jun, CD14* and *HDAC7* mRNAs as well as *iNOS* and *IL-1β* mRNA levels were significantly increased in WT cells expressing MEF2D-S444A compared to WT cells expressing MEF2D (**Fig. 5E**). Altogether, these data indicate that phosphorylation at Ser444 represses MEF2D transcriptional activity.

#### Conclusion

Our results determine that p38γ/δKIKO mice could be an excellent tool to elucidate the *in vivo* roles of p38γ/p38δ, as their use helped us to establish specific roles for p38γ/p38δ in immune and inflammatory responses independent of TPL2. Mice and cells expressing inactive kinases are powerful tools: 1) in the study of the role of kinase binding interactions that are independent of the kinase catalytic activity, 2) to also avoid unspecific side effects caused by the treatment with small molecule inhibitors, and 3) to overcome the problems of protein compensation, due to kinase functional redundancy, inherent in assessing kinase function using knock-out mice or cells.

p38MAPK are a clear example of highly related family members with functional redundancies, which may account for the difficulty on finding altered phenotypes in the p38γ- or p38δ-null mice. This highlights the need for using mice expressing inactive kinases in combination with knock-out mice in the study of the role of these kinases. Here we demonstrate with the use of p38γ/δKIKO mice, that p38γ/p38δ are implicated in the immune response modulating the expression of inflammatory genes and the phosphorylation of regulatory proteins such as the transcription factor MEF2D.

Research using p38γ/δKIKO mice, or alternatively a mouse expressing both inactive p38γ and p38δ, will be of great help to investigate which inflammatory diseases are dependent on p38γ/p38δ catalytic activity. This aspect is very important for the development of p38γ/p38δ inhibitors to treat inflammatory diseases and also cancer.

## Materials and Methods

### Generation and genotyping of p38γ^D171A/D171A^/p38δ^-/-^ (p38γ/δKIKO) mice

p38γ/δKIKO mouse were produced by crossing p38γ^D171A/D171A^ and p38δ^-/-^ mice and genotype was confirmed by PCR as described elsewhere (Sabio *et al*., 2005; Sabio *et al*., 2010). All mice were housed in specific pathogen-free conditions in the CNB-CSIC animal house. Animal procedures were performed in accordance with national and EU guidelines, with the approval of the Centro Nacional de Biotecnología Animal Ethics Committee, CSIC and Comunidad de Madrid (Reference: PROEX 316/15 and PROEX 071/19).

### Antibody

The description of all antibodies and dilution used in this study is provided in the **Table EV4**.

### Cell culture, stimulation, transfection and lysis

Mouse embryonic fibroblasts (MEF) were cultured as described previously (Sabio *et al*., 2005). MEF were incubated in DMEM overnight in the absence of serum before stimulation with 0.5 M sorbitol, then lysed in lysis buffer ([50 mM Tris-HCl (pH 7.5), 1 mM EGTA, 1 mM EDTA, 50 mM sodium fluoride, 10 mM sodium β-glycerophosphate, 5 mM pyrophosphate, 0.27 M sucrose, 1% (vol/vol) Triton X-100] plus 0.1% (vol/vol) 2-mercaptoethanol, 0.1 mM phenylmethylsulfonyl fluoride, 1 mM benzamidine and 1 mM sodium orthovanadate). Lysates were centrifuged at 20,800 *g* for 15 min at 4°C, the supernatants removed, quick frozen in liquid nitrogen and stored at -80°C until used. Transfection of MEF was carried out using poly(ethylenimine), each 6 cm dish of cells was transfected with 2-12 μg of poly(ethylenimine) (Polysciences, Eppelheim, Germany) and 1-6 μg of plasmid DNA, as described previously (Iñesta-Vaquera *et al*., 2009). Cells were lysed 24-36 h after transfection.

Bone marrow derived macrophages (BMDM) were isolated from adult mouse femur as described elsewhere (Risco *et al*., 2012). BMDMs were stimulated in 0.1-1% serum with 100 ng/ml LPS (Sigma-Aldrich) and lysed.

Peritoneal macrophages were isolated by peritoneal lavage with sterile and cold PBS, 3 days after peritoneal injection with 2 ml of 3% (w/v) thioglycolate. Cells were plated in RPMI 1640 medium supplemented with 10% (v/v) FBS, 2 mM glutamine, 100 U/ml penicillin and 100 μg/ml streptomycin for 5 h. To reduce basal phosphorylation, medium was changed to fresh medium containing 1% (v/v) FBS for 12 h before stimulation with LPS (100 ng/ml) in fresh medium and then lysed.

For mRNA expression analysis, cells were lysed with NZYol (NZYtech) and RNA extracted using a standard protocol with chloroform-isopropanol-ethanol.

### Immunoblot analysis

Protein samples were resolved in SDS-PAGE and transferred to nitrocellulose membranes, blocked (30 min) in TBST buffer (50 mM Tris/HCl pH 7.5, 0.15 M NaCl, 0.05% (v/v) Tween) with 5% (w/v) dry milk, then incubated in TBST buffer with 5% (w/v) dry milk and 0.5-1 μg/ml antibody (2 h, room temperature (RT) or overnight, 4°C). Protein was detected using fluorescently labelled secondary antibodies (Invitrogen) and the Licor Odyssey infrared imaging system.

### Immunoprecipitation

Extracts from MEF were incubated with 2 μg anti-hDlg or 2 μg anti-p38γ antibody coupled to protein-G-Sepharose. After incubation for 2 h at 4°C, the captured proteins were centrifuged at 20,800 *g* for 1 min, the supernatants discarded and the beads washed twice in lysis buffer containing 0.5 M NaCl, then twice with lysis buffer alone.

### RNA Sequencing

RNA was isolated from 1 × 106 BMDM from WT or p38γ/δKIKO male mice using the RNeasy kit (QIAGEN). Biological replicate libraries were prepared using the TruSeq RNA library prep kit (Illumina) and were single-end sequenced on the Illumina HiSeq 2500 platform as described in (Blair *et al*., 2022).

### Gene expression analysis

cDNA for real-time quantitative PCR (qPCR) was generated from total RNA using the High Capacity cDNA Reverse Transcription Kit (Applied Biosystems). Real-time qPCR reactions were performed in triplicate as described (Risco *et al*., 2012) in MicroAmp Optical 384-well plates (Applied Biosystems). PCR reactions were carried out in an ABI PRISM 7900HT (Applied Biosystems) and SDS v2.2 software was used to analyse results by the Comparative Ct Method (ΔΔCt). X-fold change in mRNA expression was quantified relative to non-stimulated wild-type cells, and β-actin mRNA was used as control. Primers sequences are listed in the **Table EV5**.

### Cytokine array

Macrophages’ culture media were collected 6 h after LPS stimulation and kept at -20ºC. 450 μl of cell culture media was treated and incubated with the Proteome profiler mouse cytokine array membrane (R&D Systems) according to the manufacturer’s instructions. Proteins were visualised using chemiluminescent detection reagent and signal intensity was determined using ImageJ software.

### Phosphoproteomic analysis

Following determination of protein by Bradford, 0.5 mg peritoneal macrophage protein lysate (samples were in triplicate) were reduced and alkylated in 50 mM ammonium bicarbonate, 10 mM TCEP (Tris(2-carboxyethyl)phosphine hydrochloride; Thermo Fisher Scientific), 55 mM chloroacetamide(Sigma-Aldrich) for 1 h at 37°C. Samples were digested with 10 μg of trypsin for 8 h at 37°C and dried down in a speed-vac. For phosphopeptide enrichment on titanium oxide (TiO_2_) columns (GL Sciences), tryptic peptides were dissolved in 0.25 M lactic acid/3% trifluoracetic acid (TFA)/70% acetonitrile (ACN) and centrifuged at 13,000 rpm for 5 min at room temperature. The supernatant was loaded on the TiO_2_ microcolumn previously equilibrated with 100 μl of 3% TFA/70% ACN. Each microcolumn was washed with 100 μl of lactic acid solution and 200 μl of 3% TFA/70% ACN. Phosphopeptides were eluted with 200 μl of 1% NH_4_OH pH 10 in water and acidified with 7 μl of TFA. Eluates were desalted using Oasis HLB cartridges, dried down and resolubilized in 5% ACN-0.2% formic acid (FA). The phosphopeptides were separated on a home-made reversed-phase column (150-μm i.d. by 150 mm) with a 240-min gradient from 10 to 30% ACN-0.2% FA and a 600-nl/min flow rate on a Nano-LC-Ultra-2D (Eksigent, Dublin, CA) connected to a Q-Exactive Plus (Thermo Fisher Scientific, San Jose, CA). Each full MS spectrum acquired at a resolution of 70,000 was followed by 12 tandem-MS (MS-MS) spectra on the most abundant multiply charged precursor ions. Tandem-MS experiments were performed using higher-energy collisional dissociation (HCD) at a collision energy of 25%. The data were processed using PEAKS 8.5 (Bioinformatics Solutions, Waterloo, ON) and a Uniprot mouse database. Mass tolerances on precursor and fragment ions were 10 ppm and 0.01 Da, respectively. Variable selected posttranslational modifications were carbamidomethyl (C), oxidation (M), deamidation (NQ), acetyl (N-term) and phosphorylation (STY).

### p38MAPK in vitro kinase assay

Kinase assay were performed in 30 μl final phosphorylation reaction mixture containing GST-MEF2D (0.4 μM, 1 μg), active p38MAPK (0.5 U/ml) and 50 mM Tris-HCl pH 7.5, 0.1 mM EGTA, 10 mM MgCl_2_ and 0.1 mM [γ^32^P]ATP (specific activity: ∼3 ×10^6^ cpms). The reactions were carried out at 30ºC for 60 min and terminated by adding 4 x SDS-PAGE sample buffer containing 1% (v/v) 2-mercaptoethanol. Myelin basic protein (MBP) phosphorylation was performed under the same conditions, but reaction was stopped by spotting the phosphorylation reaction mixture onto P81 filtermats, washed four times in 75 mM phosphoric acid to remove ATP, washed once in acetone, dried and counted for radioactivity.

### In-gel sample digestion for LC/MS–MS

Purified recombinant GST-MEF2D was phosphorylated using active p38α or p38δ (0.5 U/ml) in a buffer containing 50 mM Tris-HCl pH 7.5, 0.1 mM EGTA, 10 mM MgCl_2_ and 0.1 mM ATP for 1 h at 30ºC. The reaction was stopped by addition of 4 x SDS-PAGE sample buffer containing 1% (v/v) 2-mercaptoethanol and samples were subjected to electrophoresis on 10% SDS-PAGE gels. The gels were Coomassie stained and bands corresponding to MEF2D were excised, cut into pieces (∼1mm^2^) and manually processed for in-gel digestion. The digestion protocol used was based on Schevchenko et al. (Shevchenko *et al*., 1996) with minor variations: gel pieces were washed with 50 mM ammonium bicarbonate and then with acetonitrile (ACN), prior to reduction (10 mM DTT in 25 mM ammonium bicarbonate solution) and alkylation (55 mM iodoacetamide in 50 mM ammonium bicarbonate solution). Gel pieces were then washed with 50 mM ammonium bicarbonate, with ACN (5 min each), and then were dried under a stream of nitrogen. Pierce MS-grade trypsin (Thermo-Fisher Scientific) was added at a final concentration of 20 ng/μl in 50 mM ammonium bicarbonate solution, for overnight at 37°C. Peptides were recovered in 50% ACN / 1% formic acid (FA), dried in speed-Vac and kept at -20°C until phosphopeptide enrichment. Phosphopeptide enrichment procedure concatenated two in-house packed microcolumns, Immobilized Metal Affinity Chromatography (IMAC) and Oligo R3 polymeric reversed-phase that provided selective purification and sample desalting prior to LC-MS/MS analysis, and was performed as previously reported (Navajas *et al*., 2011).

### Protein identification by tandem mass spectrometry (LC–MS/MS Exploris 240)

The peptide samples were analysed on a nano liquid chromatography system (Ultimate 3000 nano HPLC system, Thermo Fisher Scientific) coupled to an Orbitrap Exploris 240 mass spectrometer (Thermo Fisher Scientific). Samples (5 μl) were injected on a C18 PepMap trap column (5 μm, 100 μm I.D. x 2 cm, Thermo Scientific) at 10 μl/min, in 0.1% formic acid in water, and the trap column was switched on-line to a C18 PepMap Easy-spray analytical column (2 μm, 100 Å, 75 μm I.D. x 50 cm, Thermo Scientific). Equilibration was done in mobile phase A (0.1% formic acid in water), and peptide elution was achieved in a 30 min gradient from 4% - 50% B (0.1% formic acid in 80% acetonitrile) at 250 nl/min. Data acquisition was performed using a data-dependent top-15 method, in full scan positive mode (range of 350 to 1200 m/z). Survey scans were acquired at a resolution of 60,000 at m/z 200, with Normalized Automatic Gain Control (AGC) target of 300 % and a maximum injection time (IT) of 45 ms. The top 15 most intense ions from each MS1 scan were selected and fragmented by Higher-energy collisional dissociation (HCD) of 28. Resolution for HCD spectra was set to 15,000 at m/z 200, with AGC target of 75 % and maximum ion injection time of 80 ms. Precursor ions with single, unassigned, or six and higher charge states from fragmentation selection were excluded.

MS and MS/MS raw data were translated to mascot general file (mgf) format using Proteome Discoverer (PD) version 2.4 (Thermo Fisher Scientific) and searched using an in-house Mascot Server v. 2.7 (Matrix Science, London, U.K.) against a human database (reference proteome from Uniprot Knowledgebase). Search parameters considered fixed carbamidomethyl modification of cysteine, and the following variable modifications: methionine oxidation, phosphorylation of serine/threonine/tyrosine, deamidation of asparagine/glutamine and acetylation of the protein N-terminus. Peptide mass tolerance was set to 10 ppm and 0.02 Da, in MS and MS/MS mode, respectively, and 3 missed cleavages were allowed. The Mascot confidence interval for protein identification was set to ≥ 95% (*p* < 0.05) and only peptides with a significant individual ion score of at least 30 were considered. If there were two or more residues susceptible to phosphorylation in such a way that an alternative fragment ion assignment was possible, Mascot provided information on the probability percentage that the considered residue is phosphorylated (site analysis), so that percentages above 80 % usually correspond to a highly reliable assignment of the phosphorylation site. In addition, manual validation of the identified phosphopeptides was performed.

### LPS/D-Gal-Induced Endotoxic Shock

A combination of *E. coli* 0111:B LPS (50 μg/kg body weight) (Sigma-Aldrich) and D-Gal (1 g/kg body weight) (Sigma-Aldrich) was simultaneously injected intraperitoneally in 12-week-old WT, p38γ/δ^−/−^ and p38γ/δKIKO mice of both sexes (Risco *et al*., 2012). Plasma samples were collected 2 h or 6 h after LPS/D-Gal injection for cytokine or transaminase analysis, respectively. For TUNEL and liver haemorrhage analysis livers were collected 6 h after LPS/D-Gal injection.

### Serum analysis

Cytokine concentrations in mouse serum samples were measured using the Luminex-based MilliPlex Mouse cytokine/chemokine immunoassay and the Luminex-based Bio-Plex Mouse Grp I Cytokine 23-Plex Panel (Bio-Rad). Transaminase activities were measured using the ALT and AST Reagent Kit (Biosystems Reagents) with a Ultrospec 3100 pro UV/Visible Spectrophotometer (Amersham Biosciences).

### Histologic analysis

Histological analysis was performed on Haematoxylin and eosin (H&E)-stained sections of formalin-fixed, paraffin-embedded liver.

Apoptosis was determined by IHC in deparaffinized sections using TUNEL (terminal deoxynucleotidyl transferase–mediated dUTP nick end labelling) staining. TUNEL-positive cells were counted on slides from 25 random sections per mouse. Slides were mounted for fluorescence with Hoechst-containing mounting medium (Sigma) and analysed with a TCS SP5 Microscope (Leica).

### Infection by *C. albicans*

*C. albicans* (strain SC5314) was grown on YPD Agar (Sigma Y1500) plates at 30°C for 48 h. 8-12-week-old female mice were infected intravenously with 1 × 10^5^ colony-forming units (CFU) of *C. albicans*. Kidney fungal burden was determined at 3 days post-infection by plating the kidney homogenates in serial dilutions on YPD agar plates (Alsina-Beauchamp et al., 2018).

### Generation of MEF2D S444A mutant

MEF2D S444A (MEF2D^S444A^) mutant was generated using the NZY Supreme Mutagenesis kit following the manufacturer’s instructions. The primer used were: *Forward*: 5’ TCA CGG CTT GGG GCC ACC GGT TCT GAC 3’ *Revers*: 5’ GTC AGA ACC GGT GGC CCC AAG CCG TGA 3’

### Luciferase reporter assay

WT, p38γ/δ^-/-^ or p38γ/δKIKO mouse embryonic fibroblasts (MEFs) were seeded in p24 plates (25,000 cells per well) and incubated overnight. The following days, 100 ng (per well) of Flag-MEF2D or Flag-MEF2D^S444A^ were co-transfected with 500 ng (per well) of luciferase plasmids MEF-luciferase and renilla, using Lipofectamine 2000 (ThermoFisher) according to the manufacturer’s instructions. After 24h, cells were lysed using Passive Lysis Buffer (Promega). When p38MAPK inhibitors were used, cells were treated with the indicated concentrations of BIRB796 or SB203580 6 h prior to cell lysis. Cell lysates were used to measure Luciferase activities with the Dual-Luciferase Reporter Assay System (Promega) as indicated by the manufacturer’s instructions, using an Infinite M200 luminometer (TECAN). Renilla and Firefly luciferase intensities were used to calculate luciferase activity.

### Cell recruitment analysis by flow cytometry

Leukocyte infiltration in the kidney was analysed by flow cytometry as described (Barrio *et al*., 2020). Briefly, flow cytometry analysis was performed on cell suspensions from kidney homogenates that were digested with 0.2 mg/ml liberase (Roche) and 0.1 mg/ml DNase I (Roche) for 20 min at 37°C on a shaking platform and filtered through 40 μm cell strainers (Falcon). Cells were stained with combinations of fluorescent-labelled antibodies against the cell surface markers CD45, Ly6G and F4/80 and analysed in a Cytomics FC500 flow cytometer (Beckman Coulter). Profiles were analysed with Kaluza software (Beckman Coulter); leukocytes were gated as CD45+cells as described. Cell subpopulations in bone marrow and spleen were analysed as described in (Barrio *et al*., 2020).

### Statistical analysis

*In vitro* experiments have been performed at least twice with three independent replicates per experiment. For the analysis of mouse survival, production of inflammatory molecules and transaminases, the groups’ size was established according to the Spanish ethical legislation for animal experiments. At least 5 mice per group were used. Differences in mouse survival were analysed by two-way ANOVA using GraphPad Prism software. Other data were analysed using Student’s t-test. In all cases, *p*-values < 0.05 were considered significant. Data are shown as mean ± SEM.

## Acknowledgments

We thank V. Marquez (CNB-CSIC) for technical support; the antibody purification teams (Division of Signal Transduction Therapy, University of Dundee), coordinated by H. McLauchlan and J. Hastie, for generation and purification of antibodies. We thank the Histology Facility at CNB-CSIC for the histological processing of biological samples. This research was funded by the MCIN/AEI/10.13039/501100011033 (PID2019-108349RB-100 and SAF2016-79792R) to AC and JJSE; and by a grant from the Francis Crick Institute, which receives its core funding from Cancer Research UK (FC001103), the UK Medical Research Council (FC001103) and the Wellcome Trust (FC001103) to SCL. AE, PF and DG-R receive MCIN FPI fellowships, ED-M MEFP FPU fellowship and J M-G a MEFP FPU and a MCIN-Residencia de Estudiantes fellowship. MEF2D phosphorylation sites were i dentified in the proteomics facility of Centro Nacional de Biotecnología (CSIC). PF, DG-R, ED-M and J M-G are in the PhD Programme in Molecular Bioscience, Doctoral School, Universidad Autónoma de Madrid, 28049-Madrid, Spain.

## Author contributions

A.C. designed research with A.E., E.D-M and J.J.S-E.; A.E., E.D-M, P.F, D.G-R, A.R. and J.M-G. performed experiments; E.B. performed the phosphoproteomic analysis; S.C.L and M.P. performed RNA-Seq and the statistical analysis of RNA-Seq. N.S. and S.M.J provided reagents for luciferase assays and to interpret the data; A.C wrote the manuscript with the help of all other authors.

## References

Alsina-Beauchamp D, Escos A, Fajardo P, Gonzalez-Romero D, Diaz-Mora E, Risco A, Martin-Serrano MA, Del Fresno C, Dominguez-Andres J, Aparicio N et al. (2018) Myeloid cell deficiency of p38γ/p38δ protects against candidiasis and regulates antifungal immunity. EMBO Mol Med 10: e8485.

Arthur JS, Ley SC (2013) Mitogen-activated protein kinases in innate immunity. Nat Rev Immunol 13: 679–692.

Barrio L, Roman-Garcia S, Diaz-Mora E, Risco E, Jimenez-Saiz R, Carrascoo YR, Cuenda A (2020) B Cell Development and T-Dependent Antibody Response Are Regulated by p38γ and p38δ. Front Cell Dev Biol 8: 189.

Blair L, Pattison MJ, Chakravarty P, Papoutsopoulou S, Bakiri L, Wagner EF, Smale S, Ley SC (2022) TPL-2 Inhibits IFN-β Expression via an ERK1/2-TCF-FOS Axis in TLR4-Stimulated Macrophages. J Immunol 208: 164–178.

Chan SF, Sances S, Brill LM, Okamoto S-I, Zaidi R, McKercher SR, Akhtar MW, Nakanishi N, Lipton SA (2014) ATM-dependent phosphorylation of MEF2D promotes neuronal survival after DNA damage. J Neurosci 34: 4640–4653.

Cohen P (2009) Targeting protein kinases for the development of anti-inflammatory drugs. Curr Opin Cell Biol 21: 317–324.

Cuenda A, Rousseau S (2007) p38 MAP-kinases pathway regulation, function and role in human diseases. Biochim Biophys Acta 1773: 1358–1375.

Cuenda A, Sanz-Ezquerro JJ (2017) p38gamma and p38delta: From Spectators to Key Physiological Players. Trends Biochem Sci 42:431–442.

Del Reino P, Alsina-Beauchamp D, Escos A, Cerezo-Guisado MI, Risco A, Aparicio N, Zur R, Fernandez-Estevez M, Collantes E, Montans J et al. (2014) Pro-oncogenic role of alternative p38 mitogen-activated protein kinases p38γ and p38δ, linking inflammation and cancer in colitis-associated colon cancer. Cancer Res 74: 6150–6160.

Dumitru CD, Ceci JD, Tsatsanis C, Kontoyiannis D, Stamatakis K, Lin JH, Patriotis C, Jenkins NA, Copeland NG, Kollias G et al. (2000) TNF-alpha induction by LPS is regulated posttranscriptionally via a Tpl2/ERK-dependent pathway. Cell 103: 1071–1083.

Escós A, Martin-Gomez J, Gonzalez-Romero D, Diaz-Mora E, Francisco-Velilla R, Santiago C, Cuezva JM, Dominguez-Zorita S, Martinez-Salas E, Sonenberg N et al. (2022) TPL2 kinase expression is regulated by the p38γ/p38δ-dependent association of aconitase-1 with TPL2 mRNA. Proc Natl Acad Sci U S A 119: e2204752119.

Gaestel M, Kotlyarov A, Kracht M (2009) Targeting innate immunity protein kinase signalling in inflammation. Nat Rev Drug Discov 8: 480–499.

Gantke T, Sriskantharajah S, Ley SC (2011) Regulation and function of TPL-2, an IκB kinase-regulated MAP kinase kinase kinase. Cell Res 21: 131–145.

Han TH, Prywes R (1995) Regulatory role of MEF2D in serum induction of the c-jun promoter. Mol Cell Biol 15: 2907–2915.

Han J, Wu J, Silke J (2020) An overview of mammalian p38 mitogen-activated protein kinases, central regulators of cell stress and receptor signaling. F1000Res 9.

Iñesta-Vaquera FA, Centeno F, Del Reino P, Sabio G, Peggie M, Cuenda A (2009) Proteolysis of the tumour suppressor hDlg in response to osmotic stress is mediated by caspases and independent of phosphorylation. FEBS J 276: 387–400.

Kato Y, Zhao M, Morikawa A, Sugiyama T, Chakravortty D, Koide N, Yoshida T, Tapping RI, Yang Y, Yokochi T et al. (2000) Big mitogen-activated kinase regulates multiple members of the MEF2 protein family. J Biol Chem 275: 18534–18540.

Ke K, Shen J, Song Y, Cao M, Lu H, Liu C, Shen J, Li A, Huang J, Ni H (2015) CDK5 contributes to neuronal apoptosis via promoting MEF2D phosphorylation in rat model of intracerebral hemorrhage. J Mol Neurosci 56: 48–59.

Koliaraki V, Roulis M, G. Kollias G (2012) Tpl2 regulates intestinal myofibroblast HGF release to suppress colitis-associated tumorigenesis. J Clin Invest 122: 4231–4242.

Kuma Y, Sabio G, Bain J, Shpiro N, Marquez R, Cuenda A (2005) BIRB796 inhibits all p38 MAPK isoforms in vitro and in vivo. J Biol Chem 280: 19472–19479.

Lee MS, Kim YJ (2007) Signaling pathways downstream of pattern-recognition receptors and their cross talk. Annu Rev Biochem 76: 447–480.

Lu F, Wang R, Xia L, Nie T, Gao F, Yang S, Huang L, Shao K, Liu J, Yang Q (2021) Regulation of IFN-Is by MEF2D Promotes Inflammatory Homeostasis in Microglia. J Inflamm Res 14: 2851–2863.

McKinsey TA, Zhang CL, Olson EN (2002) MEF2: a calcium-dependent regulator of cell division, differentiation and death. Trends Biochem Sci 27: 40–47.

Navajas R, Paradela A, Albar JP (2011) Immobilized metal affinity chromatography/reversed-phase enrichment of phosphopeptides and analysis by CID/ETD tandem mass spectrometry. Methods Mol Biol 681: 337–348.

Park SY, Shin HM, Han TH (2002) Synergistic interaction of MEF2D and Sp1 in activation of the CD14 promoter. Mol Immunol 39: 25–30.

Pattison MJ, Naik RJ, Reyskens KMSE, Arthur JSC (2020) Loss of Mef2D function enhances TLR induced IL-10 production in macrophages. Biosci Rep 40: BSR20201859.

Risco A, Del Fresno C, Mambol A, Alsina-Beauchamp D, MacKenzie KF, Tang H-T, Barber DF, Morcelle C, Arthur JS, Ley SC et al. (2012) p38gamma and p38delta kinases regulate the Toll-like receptor 4 (TLR4)-induced cytokine production by controlling ERK1/2 protein kinase pathway activation. Proc Natl Acad Sci U S A 109: 11200–11205.

Sabio G, Arthur JS, Kuma Y, Peggie M, Carr J, Murray-Tait V, Centeno F, Goedert M, Morrice NA, Cuenda A (2005) p38gamma regulates the localisation of SAP97 in the cytoskeleton by modulating its interaction with GKAP. EMBO J 24: 1134–1145.

Sabio G, Cerezo-Guisado MI, Del Reino P, Iñesta-Vaquera Fa, Rousseau S, Arthur JS, Campbell DG, Centeno F, Cuenda A (2010) p38gamma regulates interaction of nuclear PSF and RNA with the tumour-suppressor hDlg in response to osmotic shock. J Cell Sci 123: 2596–2604.

Shevchenko A, Wilm M, Vorm O, Mann M (1996) Mass spectrometric sequencing of proteins silver-stained polyacrylamide gels. Anal Chem 68: 850–858.

Takeuchi O, Akira S (2010) Pattern recognition receptors and inflammation. Cell 140: 805–820.

Tang X, Wang X, Gong X, Tong M, Park D, Xia Z, Mao Z (2005) Cyclin-dependent kinase 5 mediates neurotoxin-induced degradation of the transcription factor myocyte enhancer factor 2. J Neurosci 25: 4823–4834.

Wang P, Zhao J, Sun X (2021) DYRK1A phosphorylates MEF2D and decreases its transcriptional activity. J Cell Mol Med 25: 6082–6093.

Yang S, Gao L, Lu F, Wang B, Gao F, Zhu G, Cai Z, Lai J, Yang Q (2015) Transcription factor myocyte enhancer factor 2D regulates interleukin-10 production in microglia to protect neuronal cells from inflammation-induced death. J Neuroinflammation 12: 33.

