## Supplementary figures and tables for "p38γ and p38δ modulate innate immune response by regulating MEF2D activation"

### Expanded View Figure Legends

**Figure EV1. Characterization of the p38γ/δKIKO mouse.**

**(A)** Samples of genomic DNA from WT, p38δ$^{-/-}$, p38γ$^{171A/171A}$ and p38γ/δKIKO purified from tail biopsy were used as templates for PCR as described in methods.

(**B**) WT and p38γ/δKIKO mouse embryonic fibroblasts (MEFs) were exposed to 0.5 M sorbitol (15 min), activation and expression of MAPK was examined in immunoblotted lysates (30 µg) using phospho-specific and specific antibodies for each indicated protein.

(**C**) MEFs were treated as in (B); endogenous p38γ immunoprecipitated with anti-p38γ from 2 mg cell lysate and immunoblotted with the p38 phospho-specific antibody, which recognises all phosphorylated p38 isoforms.

(**D**) Endogenous hDlg was immunoprecipitated with anti-hDlg from 1 mg MEF lysate (treated as in (B)). Pellets were immunoblotted with the indicated antibodies. Red (*) mark unspecific protein bands.

(**E**) Extracts from WT, p38γ/δ$^{-/-}$ and p38γ/δKIKO peritoneal macrophages were immunoblotted using specific antibodies for TPL2 and p38α. Representative blots of two independent experiments are shown in all panels.

**Figure EV2. Characterization of immune cell populations of the p38γ/δKIKO mouse.**

(**A**) *In vitro* BMDM development is not affected in p38γ/δKIKO mice. BMDM from WT and p38γ/δKIKO mice were stained with anti-F4/80 antibody, and analysed by flow cytometry. Results are representative plots (*n* = 3).

(**B**) BM or spleen cell suspensions were stained with the indicated antibodies and analysed by flow cytometry. Representative flow cytometry profiles and analysis are shown.

(**C**) Frequency and (**D**) total cell number of leukocytes (CD45$^+$), myeloid cells (CD11b$^+$), neutrophils (Ly6G$^+$) and macrophages (F4/80$^+$) in adult mouse BM or spleen. Percentage of myeloid cell population was determined relative to $CD45^+$ cell and percentages of neutrophils and macrophages were determined relative to $CD45^+$ $CD11b^+$ cell.

(**E**) Total cell number in BM or spleen of the indicated genotypes.

(**F**) Weight of the spleen of the indicated genotypes. Each dot in C-F represents a single mouse (*n*= 4-5). ns, not significant; * $p \leq 0.05$.

**Figure EV3. RNA-sequencing analysis and cytokine production in LPS-stimulated WT and p38γ/δKIKO macrophages.**

(**A**) Venn diagrams showing overlaps of genes upregulated or downregulated (-1.5< logFC <1.5, $p \leq 0.05$) in p38γ/δKIKO macrophages over the time course of LPS stimulation, compared to WT.

(**B**) Table showing GO biological processes genes.

(**C**) BMDM from WT and p38γ/δKIKO mice were exposed to LPS (100 ng/ml) for 6 h. Cell culture media (450 µl) from WT and p38γ/δKIKO macrophages were incubated with the Proteome profiler mouse cytokine array membrane according to the manufacturer's instructions. Proteins were visualised using chemiluminescent detection reagent for 30 sec. and then performing short exposure (~ 0.5 min) and long exposure (~ 5 min). Representative membranes are shown.

(**D**) Corresponding quantification by determining pixel densities on the film using ImageJ software. ns, not significant; * $p \leq 0.05$; ** $p \leq 0.01$.

**Figure EV4. Identification of proteins phosphorylated by p38γ/p38δ.**

(**A**) Peritoneal macrophages from WT, p38γ/$\delta^{-/-}$ and p38γ/δKIKO mice were exposed to LPS (100 ng/ml) for the indicated times. Cell lysates (30 µg) were immunoblotted with the indicated antibodies to active phosphorylated p38α (P-p38α) and ERK1/2 (P-ERK1/2). Total protein levels of p38α and ERK1/2 were also measured in the same lysates. Representative blots of two independent experiments with similar results are shown.

(**B**) Peritoneal macrophages and BMDM from WT, p38γ/δ$^{-/-}$ and p38γ/δKIKO mice were exposed to LPS (100 ng/ml) for the indicated times. Relative mRNA expression of indicated genes at different times was determined by qPCR and normalised to β-actin mRNA. Data show mean ± SEM ($n$ = 3-6). * $p \leq 0.05$; ** p ≤0.01, *** $p \leq 0.001$, relative to WT mice in the same conditions.

(**C**) Phosphorylated peptides from LPS-stimulated macrophages were identified. Volcano plots show the phosphoproteome statistical significance in LPS-treated macrophages between times 0 and 60. $n$= 3 per condition.

**Figure EV5. Identification of MEF2D residues phosphorylated in vitro by p38α and p38δ.**

(**A**) LC-MS/MS analysis of the *m/z* ion of the phosphorylated MEF2D peptide (amino acids and phosphopeptide sequences are indicated, where pS (red) indicates phosphorylated serine).

(**B**) The intensity of the MEF2D-derived phosphopeptides was expressed as abundance of the corresponding *m/z*.

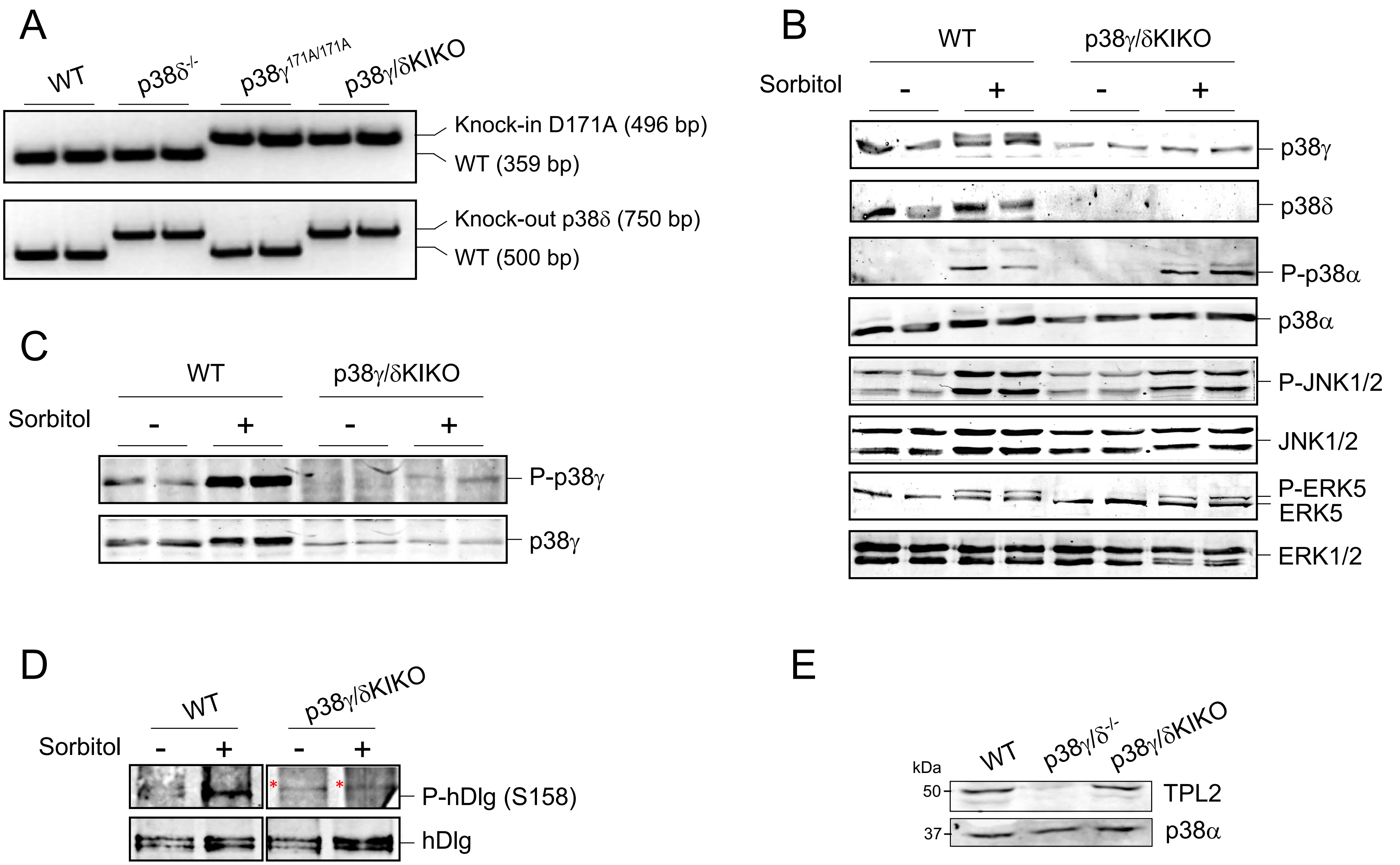
A
WT
p38δ-/-
p38γ171A/171A
p38γ/δKIKO
Knock-in D171A (496 bp)
WT (359 bp)
Knock-out p38δ (750 bp)
WT (500 bp)
B
WT
p38γ/δKIKO
Sorbitol
-
+
-
+
p38γ
p38δ
P-p38α
p38α
P-JNK1/2
JNK1/2
P-ERK5
ERK5
ERK1/2
C
WT
p38γ/δKIKO
Sorbitol
-
+
-
+
P-p38γ
p38γ
D
WT
p38γ/δKIKO
Sorbitol
-
+
-
+
P-hDlg (S158)
hDlg
E
kDa
WT
p38γ/δ-/-
p38γ/δKIKO
50
37
TPL2
p38α

Fig. EV1

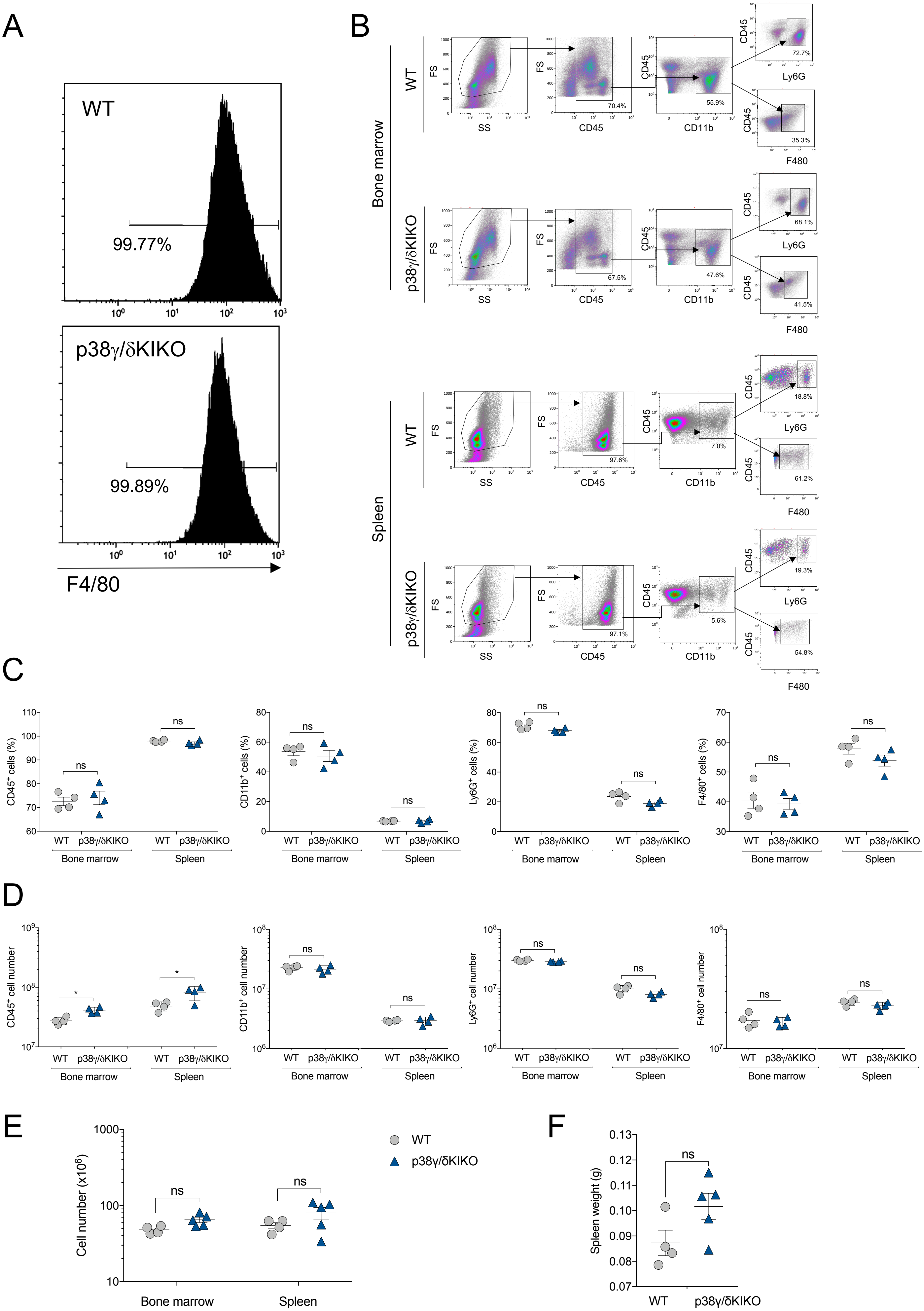
A
WT
99.77%
p38γ/δKIKO
99.89%
F4/80
B
Bone marrow
WT
p38γ/δKIKO
FS
SS
CD45
CD11b
Ly6G
F480
70.4%
55.9%
72.7%
35.3%
67.5%
47.6%
68.1%
41.5%
Spleen
97.6%
7.0%
18.8%
61.2%
97.1%
5.6%
19.3%
54.8%
C
CD45+ cells (%)
CD11b+ cells (%)
Ly6G+ cells (%)
F4/80+ cells (%)
ns
WT
p38γ/δKIKO
Bone marrow
Spleen
D
CD45+ cell number
CD11b+ cell number
Ly6G+ cell number
F4/80+ cell number
*
E
Cell number (x10^6)
F
Spleen weight (g)

Fig. EV2

A

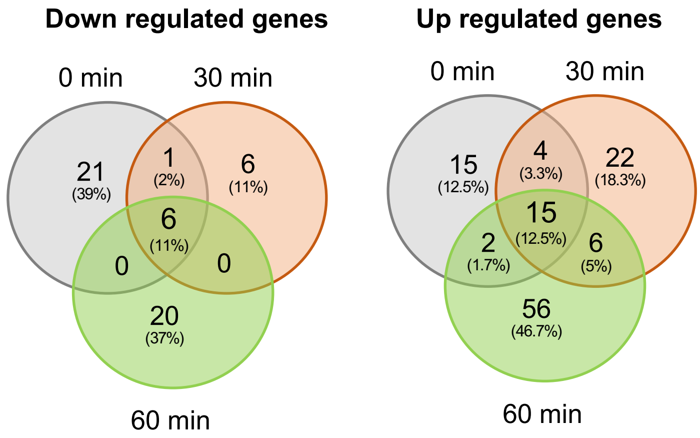


B

| Go-Biological process | Genes |
|---|---|
| Defense response to virus | IFI206, CXCL9, RSAD2, IFNA1, IFNB1, MX2, MX1, IFNA2, ACOD1, IFIT1BL1, IFIT1, IFIT3, IFIT2, CXCL10, IL6, IL12B, IFIT3B |
| Cellular response to IFN-beta | IFI208, TGTP2, IFI206, IFI205, IFNB1, CAPN2, ACOD1, IFI202B, IFIT1, IFIT3, IIGP1 |
| Defense response | CXCL10, CXCL9, TGTP2, IFNA1, IFNB1, IFNA2, ACOD1, IIGP1, CAMP |
| Inflammatory response | GBP5, CXCL9, NOS2, ACOD1, IL27, IFI202B, CXCL10, IL6, CXCL11, ITGB2L, IL1B, CCL5, CHIL1 |
| Immune response | H2-T24, CXCL9, GM43302, TNFSF15, H2-M2, H2-Q6, LIF, CXCL10, IL6, CXCL11, IL1B, CCL5, CLEC4E |
| innate immune response | TGTP2, RSAD2, TRIM30C, MX1, ACOD1, IL27, IFI202B, IFIT1, IFIT3, IIGP1, IFIT2, PTX3, CLEC4E, CAMP, CD177 |
| Negative regulation of viral genome replication | IFI206, RSAD2, IFNB1, CCL5, MX2 |
| Positive regulation of IL-8 production | IL6, NOS2, IL1B, CHIL1, CAMP |
| Positive regulation of IL-1-beta production | GBP5, IL6, IFI206, IFI205, IFI202B |
| Chemokine-mediated signaling pathway | CXCL10, CXCL9, CXCL11, CCL5 |

C

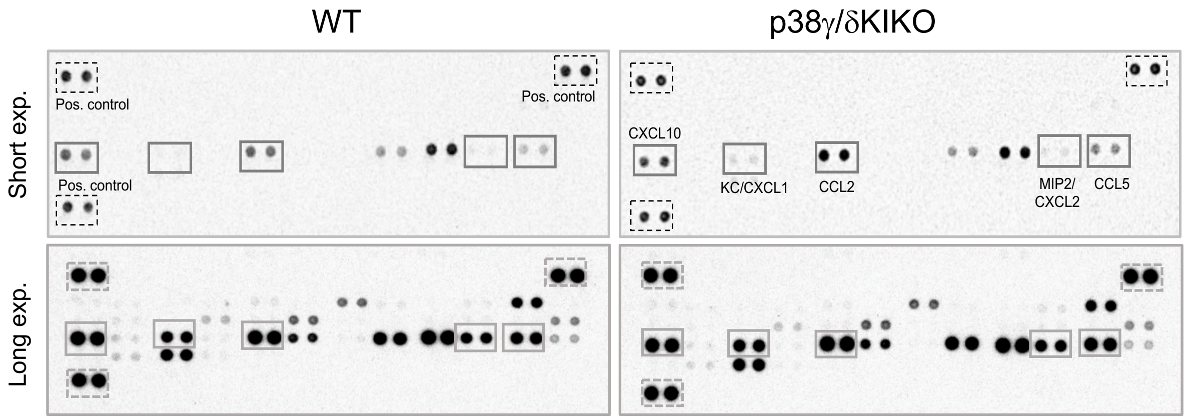


D

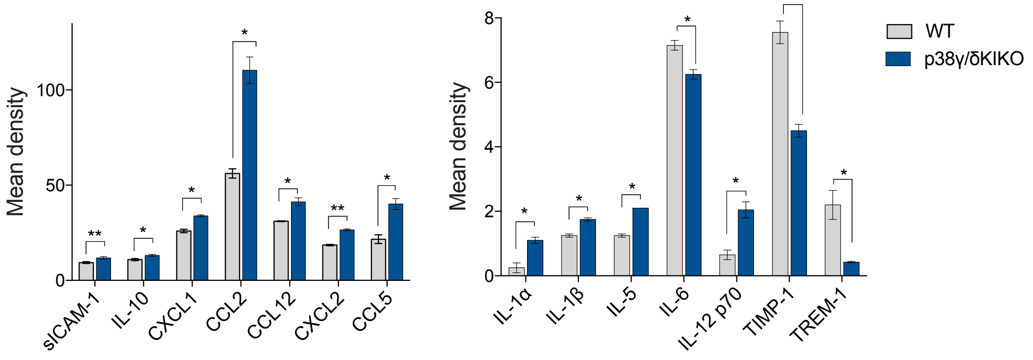


Fig. EV3

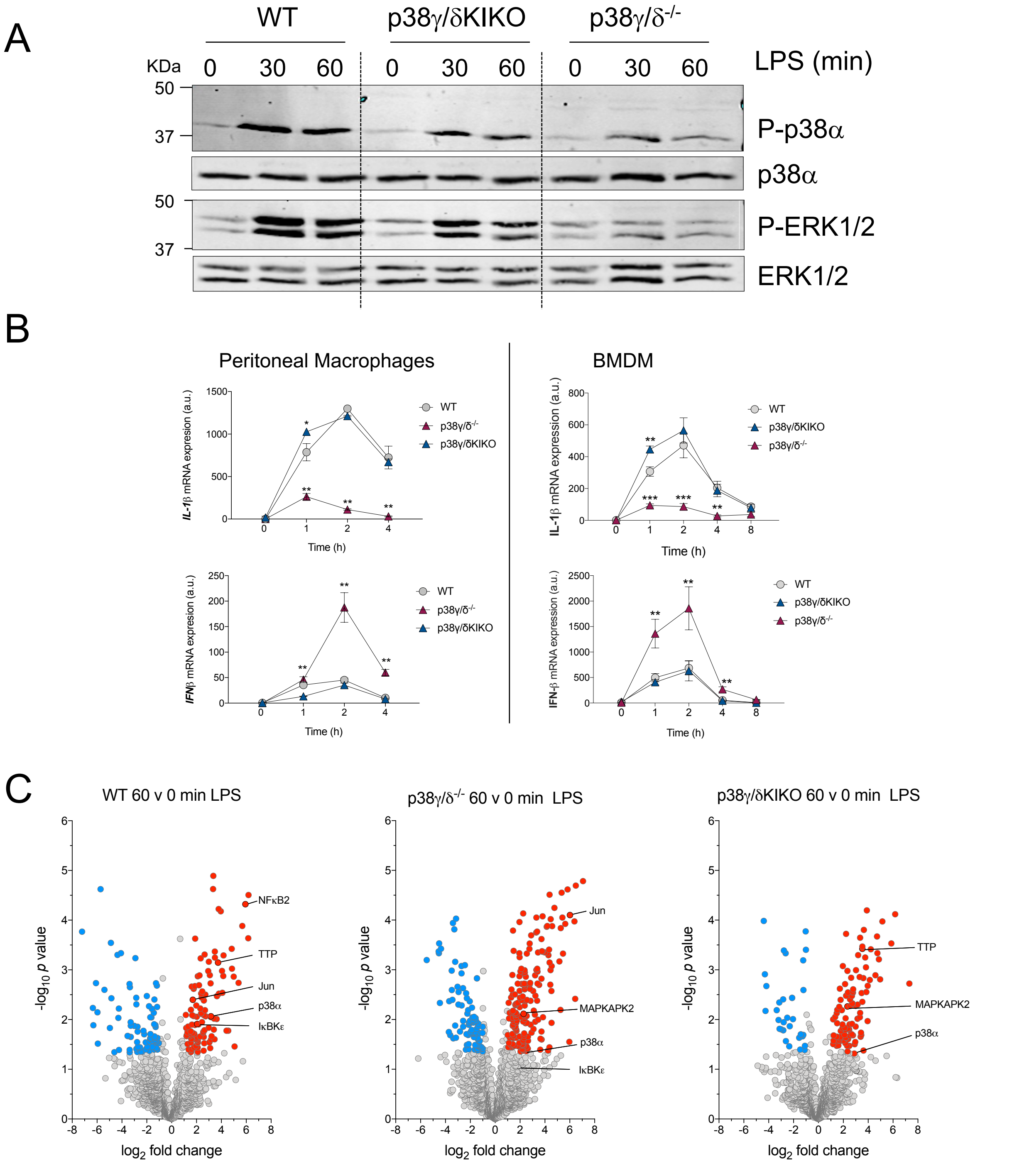
A
WT
p38γ/δKIKO
p38γ/δ-/-
KDa
0 30 60 0 30 60 0 30 60
LPS (min)
50
37
P-p38α
p38α
50
37
P-ERK1/2
ERK1/2
B
Peritoneal Macrophages
BMDM
IL-1β mRNA expresion (a.u.)
IL-1β mRNA expression (a.u.)
IFNβ mRNA expresion (a.u.)
IFN-β mRNA expression (a.u.)
Time (h)
WT
p38γ/δ-/-
p38γ/δKIKO
C
WT 60 v 0 min LPS
p38γ/δ-/- 60 v 0 min LPS
p38γ/δKIKO 60 v 0 min LPS
-log10 p value
log2 fold change
NFκB2
TTP
Jun
p38α
IκBKε
MAPKAPK2

Fig. EV4

A

B

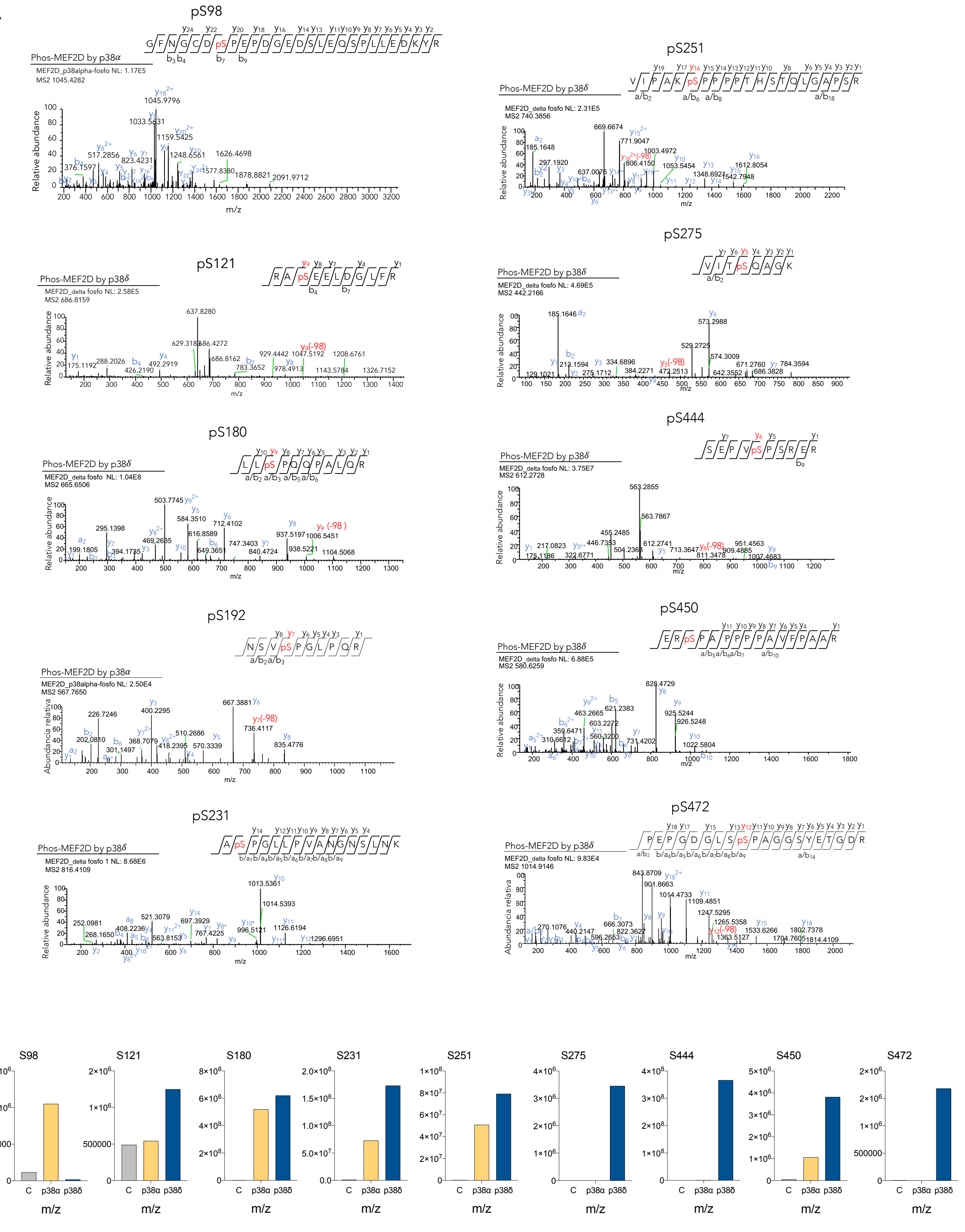
pS98
Phos-MEF2D by p38α
MEF2D_p38alpha-fosfo NL: 1.17E5
MS2 1045.4282
Relative abundance
m/z
pS251
Phos-MEF2D by p38δ
MEF2D_delta fosfo NL: 2.31E5
MS2 740.3856
pS121
Phos-MEF2D by p38δ
MEF2D_delta fosfo NL: 2.58E5
MS2 686.8159
pS275
Phos-MEF2D by p38δ
MEF2D_delta fosfo NL: 4.69E5
MS2 442.2166
pS180
Phos-MEF2D by p38δ
MEF2D_delta fosfo NL: 1.04E8
MS2 665.6506
pS444
Phos-MEF2D by p38δ
MEF2D_delta fosfo NL: 3.75E7
MS2 612.2728
pS192
Phos-MEF2D by p38α
MEF2D_p38alpha-fosfo NL: 2.50E4
MS2 567.7650
Abundancia relativa
pS450
Phos-MEF2D by p38δ
MEF2D_delta fosfo NL: 6.88E5
MS2 580.6259
pS231
Phos-MEF2D by p38δ
MEF2D_delta fosfo 1 NL: 8.68E6
MS2 816.4109
pS472
Phos-MEF2D by p38δ
MEF2D_delta fosfo NL: 9.83E4
MS2 1014.9146
S98
S121
S180
S231
S251
S275
S444
S450
S472
Abundance
C
p38α
p38δ
m/z

Fig. EV5

**Table EV1.**

**WT time 0 versus KIKO time 0 LPS**

| Row.names | baseMean | log2FoldChange | lfcSE | stat | pvalue | padj | Input.Type | MGI.Gene.Marker.ID | Symbol |
|---|---|---|---|---|---|---|---|---|---|
| ENSMUST00000182880 | 22,40859081 | -8,123658735 | 1,035687714 | -7,843733807 | 4,37E-15 | 4,15E-12 | Ensembl Transcript | 2442822 | Ifi208 |
| ENSMUST00000024839 | 91,54833171 | -2,697687384 | 0,179039548 | -15,06755026 | 2,65E-51 | 3,25E-47 | Ensembl Transcript | 104754 | Sik1 |
| ENSMUST00000082395 | 16,29535341 | -2,056406031 | 0,39013036 | -5,271074092 | 1,36E-07 | 2,17E-05 | Ensembl Transcript | 102480 | mt-Tm |
| ENSMUST00000214259 | 21,95229747 | -2,027388585 | 0,305038657 | -6,646333302 | 3,00E-11 | 1,85E-08 | Ensembl Transcript | 2177494 | Olfr111 |
| ENSMUST00000115371 | 71,75585119 | -1,976484969 | 0,147517057 | -13,39834873 | 6,18E-41 | 1,91E-37 | Ensembl Transcript | 97960 | Rnps1 |
| ENSMUST00000082423 | 49,78538356 | -1,947978767 | 0,355362933 | -5,481659984 | 4,21E-08 | 9,13E-06 | Ensembl Transcript | 102478 | mt-Tp |
| ENSMUST00000083103 | 14,00330459 | -1,794449493 | 0,458941794 | -3,909971843 | 9,23E-05 | 0,002482029 | Ensembl Transcript | 103186 | Rn7sk |
| ENSMUST00000115096 | 13,27394258 | -1,773770632 | 0,397174274 | -4,465975635 | 7,97E-06 | 0,00045968 | Ensembl Transcript | 2179061 | Plxna4 |
| ENSMUST00000185789 | 20,58630589 | -1,612427811 | 0,443872159 | -3,632640118 | 0,000280536 | 0,005443989 | Ensembl Transcript | 1926855 | Kcnq1ot1 |
| ENSMUST00000082416 | 45,06225743 | -1,590245349 | 0,361485624 | -4,399193897 | 1,09E-05 | 0,000547349 | Ensembl Transcript | 102474 | mt-Ts2 |
| ENSMUST00000174784 | 285,4341695 | -1,509984353 | 0,365909505 | -4,126660638 | 3,68E-05 | 0,001272467 | Ensembl Transcript | 5141882 | Gm20417 |
| ENSMUST00000035938 | 877,6691761 | 1,548433081 | 0,318187838 | 4,866411903 | 1,14E-06 | 0,000101636 | Ensembl Transcript | 98262 | Ccl5 |
| ENSMUST00000085102 | 163,527424 | 1,58082204 | 0,431888102 | 3,660258373 | 0,000251961 | 0,005031882 | Ensembl Transcript | 103159 | Cish |
| ENSMUST00000169733 | 167,5708517 | 1,584617841 | 0,15676246 | 10,10840123 | 5,07E-24 | 6,95E-21 | Ensembl Transcript | 1920754 | 1700071M16Rik |
| ENSMUST00000016427 | 130,7350349 | 1,624250034 | 0,407535127 | 3,985546098 | 6,73E-05 | 0,001964365 | Ensembl Transcript | 95914 | H2-M2 |
| ENSMUST00000173128 | 15,46820892 | 1,747747624 | 0,338392261 | 5,164856961 | 2,41E-07 | 3,30E-05 | Ensembl Transcript | 5011869 | Gm19684 |
| ENSMUST00000167624 | 365,2299933 | 1,809331017 | 0,12251691 | 14,76801047 | 2,36E-49 | 9,69E-46 | Ensembl Transcript | 95742 | Glo1 |
| ENSMUST00000120770 | 400,2625055 | 1,812711619 | 0,120669736 | 15,02208993 | 5,26E-51 | 3,25E-47 | Ensembl Transcript | 3649615 | Gm13443 |
| ENSMUST00000168254 | 82,45840255 | 1,938347423 | 0,555392551 | 3,490049373 | 0,000482931 | 0,007847357 | Ensembl Transcript | 3650685 | Ifit1bl1 |
| ENSMUST00000022722 | 8390,300294 | 2,037100418 | 0,502735602 | 4,052031348 | 5,08E-05 | 0,001602719 | Ensembl Transcript | 103206 | Acod1 |
| ENSMUST00000173900 | 27,63631049 | 2,060699086 | 0,224876858 | 9,163677884 | 5,02E-20 | 5,63E-17 | Ensembl Transcript | 1925060 | A930015D03Rik |
| ENSMUST00000153546 | 221,0238314 | 2,113526407 | 0,592185901 | 3,569025208 | 0,000358312 | 0,006399835 | Ensembl Transcript | 1927656 | Ms4a4c |
| ENSMUST00000172538 | 103,507649 | 2,181566246 | 0,186489099 | 11,6980899 | 1,30E-31 | 3,22E-28 | Ensembl Transcript | 2442805 | H2-T-ps |
| ENSMUST00000028881 | 13113,25707 | 2,282466805 | 0,457990838 | 4,983651674 | 6,24E-07 | 6,70E-05 | Ensembl Transcript | 96543 | Il1b |
| ENSMUST00000216328 | 12,42620999 | 2,423892105 | 0,452710356 | 5,354178615 | 8,59E-08 | 1,56E-05 | Ensembl Transcript | 2177482 | Olfr99 |
| ENSMUST00000113760 | 1321,489174 | 2,542985939 | 0,276037687 | 9,212459242 | 3,19E-20 | 3,93E-17 | Ensembl Transcript | 95958 | H2-T24 |
| ENSMUST00000024757 | 55,27353245 | 2,68040211 | 0,356798028 | 7,512379275 | 5,81E-14 | 4,78E-11 | Ensembl Transcript | 2682634 | Enpp4 |
| ENSMUST00000006956 | 31,75019108 | 2,838495388 | 0,517661513 | 5,483303896 | 4,17E-08 | 9,13E-06 | Ensembl Transcript | 98223 | Saa3 |
| ENSMUST00000163196 | 39,96033671 | 3,536198283 | 0,945711053 | 3,739195257 | 0,00018461 | 0,0040542 | Ensembl Transcript | 4937868 | Gm17041 |

**WT time 30 versus KIKO time 30 min LPS**

| Row.names | baseMean | log2FoldChange | lfcSE | stat | pvalue | padj | Input.Type | MGI.Gene.Marker.ID | Symbol |
|---|---|---|---|---|---|---|---|---|---|
| ENSMUST00000182880 | 22,40859081 | -6,784767809 | 0,982496064 | -6,905643753 | 5,00E-12 | 3,52E-09 | Ensembl Transcript | 2442822 | Ifi208 |
| ENSMUST00000068258 | 11,35607718 | -4,407958733 | 0,844456667 | -5,219875578 | 1,79E-07 | 3,24E-05 | Ensembl Transcript | 1918833 | 9130008F23Rik |
| ENSMUST00000214259 | 21,95229747 | -2,138165527 | 0,354086809 | -6,038534818 | 1,56E-09 | 4,96E-07 | Ensembl Transcript | 2177494 | Olfr111 |
| ENSMUST00000024839 | 91,54833171 | -2,117016577 | 0,195570573 | -10,82482167 | 2,63E-27 | 7,03E-24 | Ensembl Transcript | 104754 | Sik1 |
| ENSMUST00000115371 | 71,75585119 | -1,916245859 | 0,155084202 | -12,35616416 | 4,51E-35 | 2,01E-31 | Ensembl Transcript | 97960 | Rnps1 |
| ENSMUST00000108436 | 8,527011995 | -1,652660346 | 0,465706878 | -3,548713634 | 0,000387118 | 0,018703242 | Ensembl Transcript | 107610 | Zfp94 |
| ENSMUST00000201176 | 2392,059615 | 1,500372931 | 0,430865787 | 3,482228053 | 0,00049726 | 0,022790516 | Ensembl Transcript | 1352450 | Cxcl10 |
| ENSMUST00000154245 | 122,4819649 | 1,503289298 | 0,467616617 | 3,21479016 | 0,001305399 | 0,0482601 | Ensembl Transcript | 1202729 | Art3 |
| ENSMUST00000169733 | 167,5708517 | 1,548435446 | 0,174835259 | 8,856539915 | 8,25E-19 | 1,23E-15 | Ensembl Transcript | 1920754 | 1700071M16Rik |
| ENSMUST00000198803 | 478,1349011 | 1,563684263 | 0,404382317 | 3,866846292 | 0,000110252 | 0,006992895 | Ensembl Transcript | 2429943 | Gbp5 |
| ENSMUST00000210960 | 129,9756894 | 1,60795915 | 0,302748964 | 5,311196205 | 1,09E-07 | 2,11E-05 | Ensembl Transcript | 97610 | Plat |
| ENSMUST00000035938 | 877,6691761 | 1,699556266 | 0,338344067 | 5,023159654 | 5,08E-07 | 8,03E-05 | Ensembl Transcript | 98262 | Ccl5 |
| ENSMUST00000120770 | 400,2625055 | 1,770104472 | 0,130407633 | 13,57362623 | 5,74E-42 | 3,84E-38 | Ensembl Transcript | 3649615 | Gm13443 |
| ENSMUST00000059226 | 335,8751846 | 1,788113233 | 0,424191869 | 4,215340655 | 2,49E-05 | 0,002149576 | Ensembl Transcript | 101847 | Ifi205 |
| ENSMUST00000172538 | 103,507649 | 1,830742982 | 0,205472311 | 8,909925478 | 5,11E-19 | 8,54E-16 | Ensembl Transcript | 2442805 | H2-T-ps |
| ENSMUST00000062145 | 6,847632963 | 1,832235311 | 0,544098037 | 3,367472745 | 0,000758605 | 0,032332529 | Ensembl Transcript | 3045314 | 4933430I17Rik |
| ENSMUST00000109813 | 25,47144426 | 1,873971477 | 0,428470773 | 4,373627315 | 1,22E-05 | 0,001277647 | Ensembl Transcript | 88257 | Camk2b |
| ENSMUST00000167624 | 365,2299933 | 1,878134879 | 0,134009561 | 14,01493191 | 1,26E-44 | 1,69E-40 | Ensembl Transcript | 95742 | Glo1 |
| ENSMUST00000224001 | 57,4223966 | 2,059173562 | 0,303703083 | 6,780219486 | 1,20E-11 | 7,30E-09 | Ensembl Transcript | 1918576 | Treml1 |
| ENSMUST00000216328 | 12,42620999 | 2,091398284 | 0,400000915 | 5,228483747 | 1,71E-07 | 3,14E-05 | Ensembl Transcript | 2177482 | Olfr99 |
| ENSMUST00000016427 | 130,7350349 | 2,111509264 | 0,415633453 | 5,080219725 | 3,77E-07 | 6,31E-05 | Ensembl Transcript | 95914 | H2-M2 |
| ENSMUST00000006956 | 31,75019108 | 2,408262344 | 0,555378954 | 4,336250642 | 1,45E-05 | 0,001436776 | Ensembl Transcript | 98223 | Saa3 |
| ENSMUST00000113760 | 1321,489174 | 2,432291679 | 0,295090641 | 8,242523961 | 1,69E-16 | 2,26E-13 | Ensembl Transcript | 95958 | H2-T24 |
| ENSMUST00000024757 | 55,27353245 | 2,442838121 | 0,387141373 | 6,309938152 | 2,79E-10 | 1,13E-07 | Ensembl Transcript | 2682634 | Enpp4 |
| ENSMUST00000173900 | 27,63631049 | 2,450206354 | 0,323478667 | 7,57455314 | 3,60E-14 | 3,47E-11 | Ensembl Transcript | 1925060 | A930015D03Rik |
| ENSMUST00000022722 | 8390,300294 | 2,612820584 | 0,52027462 | 5,022002771 | 5,11E-07 | 8,03E-05 | Ensembl Transcript | 103206 | Acod1 |
| ENSMUST00000026845 | 33,29572672 | 2,6307359 | 0,685940456 | 3,835224872 | 0,000125449 | 0,00777264 | Ensembl Transcript | 96559 | Il6 |
| ENSMUST00000055671 | 568,2967999 | 2,924066925 | 0,515133514 | 5,676328265 | 1,38E-08 | 3,41E-06 | Ensembl Transcript | 107657 | Ifnb1 |
| ENSMUST00000173128 | 15,46820892 | 3,143408847 | 0,508071545 | 6,186941344 | 6,13E-10 | 2,16E-07 | Ensembl Transcript | 5011869 | Gm19684 |
| ENSMUST00000122808 | 7,606129609 | 4,221434903 | 1,167716159 | 3,615120737 | 0,000300208 | 0,015512275 | Ensembl Transcript | 1860203 | Cxcl11 |
| ENSMUST00000021822 | 16,66415962 | 21,33991832 | 5,446894943 | 3,917813459 | 8,94E-05 | 0,005833409 | Ensembl Transcript | 109278 | Ogn |

| WT time 60 versus KIKO time 60 min LPS | | | | | | | | | |
|---|---|---|---|---|---|---|---|---|---|
| Row.names | baseMean | log2FoldChange | lfcSE | stat | pvalue | padj | Input.Type | MGI.Gene.Marker.ID | Symbol |
| ENSMUST00000182880 | 22,40859081 | -8,168850391 | 0,956471956 | -8,540606276 | 1,34E-17 | 1,31E-14 | Ensembl Transcript | 2442822 | Ifi208 |
| ENSMUST00000024774 | 2,038620531 | -4,39210487 | 1,015648791 | -4,324432725 | 1,53E-05 | 0,001336599 | Ensembl Transcript | 1194489 | Guca1b |
| ENSMUST00000048065 | 2,300915453 | -3,58121028 | 1,089145102 | -3,288092904 | 0,001008685 | 0,024023712 | Ensembl Transcript | 1930003 | Trem3 |
| ENSMUST00000065666 | 4,54174104 | -3,3284221 | 0,838875774 | -3,967717516 | 7,26E-05 | 0,003787445 | Ensembl Transcript | 2667763 | Retnlg |
| ENSMUST00000024984 | 3,332223712 | -2,852881742 | 0,801462936 | -3,559592857 | 0,00037143 | 0,011501877 | Ensembl Transcript | 3039635 | Tmem204 |
| ENSMUST00000068258 | 11,35607718 | -2,582389892 | 0,304568641 | -8,478843664 | 2,27E-17 | 1,98E-14 | Ensembl Transcript | 1918833 | 9130008F23Rik |
| ENSMUST00000115371 | 71,75585119 | -2,301917994 | 0,139524275 | -16,49833328 | 3,77E-61 | 5,57E-57 | Ensembl Transcript | 97960 | Rnps1 |
| ENSMUST00000063956 | 6,428504955 | -2,087906669 | 0,506890295 | -4,119050396 | 3,80E-05 | 0,002381116 | Ensembl Transcript | 1916141 | Cd177 |
| ENSMUST00000214259 | 21,95229747 | -1,922656524 | 0,287190425 | -6,69470969 | 2,16E-11 | 1,28E-08 | Ensembl Transcript | 2177494 | Olfr111 |
| ENSMUST00000042318 | 3,812622197 | -1,888318374 | 0,591028346 | -3,19497091 | 0,001398449 | 0,030053944 | Ensembl Transcript | 2444310 | Fsd2 |
| ENSMUST00000000161 | 8,770942165 | -1,704791446 | 0,502404185 | -3,393266811 | 0,000690643 | 0,018023838 | Ensembl Transcript | 1277979 | Itgb2l |
| ENSMUST00000156873 | 8,632857908 | -1,616448681 | 0,401855879 | -4,022458708 | 5,76E-05 | 0,003198185 | Ensembl Transcript | 1340899 | Chil1 |
| ENSMUST00000047399 | 6,809526092 | -1,577633926 | 0,318975721 | -4,945937323 | 7,58E-07 | 0,000130154 | Ensembl Transcript | 1924846 | Adgrf1 |
| ENSMUST00000026845 | 33,29572672 | 1,533020949 | 0,337869765 | 4,537313208 | 5,70E-06 | 0,000652391 | Ensembl Transcript | 96559 | Il6 |
| ENSMUST00000160565 | 201,671234 | 1,536161407 | 0,385796098 | 3,981796126 | 6,84E-05 | 0,003608156 | Ensembl Transcript | 3646410 | Ifi206 |
| ENSMUST00000037994 | 54,79984639 | 1,554984137 | 0,370858132 | 4,192935263 | 2,75E-05 | 0,001964957 | Ensembl Transcript | 1313259 | Slfn1 |
| ENSMUST00000224046 | 15,66062993 | 1,578530576 | 0,364663818 | 4,328728268 | 1,50E-05 | 0,001326496 | Ensembl Transcript | 3698434 | A330040F15Rik |
| ENSMUST00000122808 | 7,606129609 | 1,643213563 | 0,511891865 | 3,210079463 | 0,001326983 | 0,029060353 | Ensembl Transcript | 1860203 | Cxcl11 |
| ENSMUST00000056759 | 686,9593234 | 1,643236401 | 0,360859718 | 4,553670914 | 5,27E-06 | 0,000613145 | Ensembl Transcript | 1333785 | Olfr56 |
| ENSMUST00000106828 | 41,50337212 | 1,738540596 | 0,4844649 | 3,588579059 | 0,000332485 | 0,01058574 | Ensembl Transcript | 4821257 | Trim30c |
| ENSMUST00000153546 | 221,0238314 | 1,7468236 | 0,563696487 | 3,098872605 | 0,001942585 | 0,037557497 | Ensembl Transcript | 1927656 | Ms4a4c |
| ENSMUST00000120770 | 400,2625055 | 1,756577885 | 0,115994708 | 15,14360365 | 8,35E-52 | 4,11E-48 | Ensembl Transcript | 3649615 | Gm13443 |
| ENSMUST00000019464 | 21,1948271 | 1,770115051 | 0,277462187 | 6,379662282 | 1,77E-10 | 9,71E-08 | Ensembl Transcript | 1919143 | Noxo1 |
| ENSMUST00000140417 | 4,702539516 | 1,791229642 | 0,499942077 | 3,582874344 | 0,000339834 | 0,010729083 | Ensembl Transcript | 2142489 | Ido2 |
| ENSMUST00000055671 | 568,2967999 | 1,806830465 | 0,444682597 | 4,063191314 | 4,84E-05 | 0,002894777 | Ensembl Transcript | 107657 | Ifnb1 |
| ENSMUST00000168254 | 82,45840255 | 1,809039573 | 0,531242966 | 3,405296047 | 0,000660924 | 0,017619703 | Ensembl Transcript | 3650685 | Ifit1bl1 |
| ENSMUST00000102824 | 1727,203204 | 1,816152706 | 0,411995899 | 4,408181511 | 1,04E-05 | 0,001002545 | Ensembl Transcript | 99450 | Ifit1 |
| ENSMUST00000113093 | 9,893852059 | 1,845381143 | 0,506521921 | 3,643240432 | 0,000269227 | 0,009017583 | Ensembl Transcript | 1352449 | Cxcl9 |
| ENSMUST00000046745 | 33,17350796 | 1,856237783 | 0,445863823 | 4,163239288 | 3,14E-05 | 0,002135421 | Ensembl Transcript | 3710083 | Tgtp2 |
| ENSMUST00000167624 | 365,2299933 | 1,897705996 | 0,120177846 | 15,79081382 | 3,60E-56 | 2,66E-52 | Ensembl Transcript | 95742 | Glo1 |
| ENSMUST00000119467 | 246,5659259 | 1,917585798 | 0,436708034 | 4,391001875 | 1,13E-05 | 0,001061532 | Ensembl Transcript | 3649299 | Gm12250 |
| ENSMUST00000110081 | 4,080362076 | 1,930965619 | 0,612078755 | 3,154766608 | 0,001606265 | 0,032847333 | Ensembl Transcript | 1933993 | Kcnn1 |
| ENSMUST00000066912 | 30,7544484 | 1,942572882 | 0,417316618 | 4,654913796 | 3,24E-06 | 0,000412717 | Ensembl Transcript | 1926259 | Iigp1 |
| ENSMUST00000224001 | 57,4223966 | 1,946911769 | 0,284847053 | 6,834937373 | 8,20E-12 | 5,51E-09 | Ensembl Transcript | 1918576 | Treml1 |
| ENSMUST00000162353 | 14,06225418 | 1,952647476 | 0,389704173 | 5,010589081 | 5,43E-07 | 9,90E-05 | Ensembl Transcript | 3612703 | A530040E14Rik |
| ENSMUST00000102825 | 805,0851935 | 1,966367706 | 0,476733748 | 4,124666471 | 3,71E-05 | 0,002353505 | Ensembl Transcript | 1101055 | Ifit3 |
| ENSMUST00000190447 | 116,3804944 | 1,976783902 | 0,416222938 | 4,74933917 | 2,04E-06 | 0,000295539 | Ensembl Transcript | 97244 | Mx2 |
| ENSMUST00000006956 | 31,75019108 | 1,983444692 | 0,437342214 | 4,535223516 | 5,75E-06 | 0,000653816 | Ensembl Transcript | 98223 | Saa3 |
| ENSMUST00000188363 | 16,84381175 | 1,986751094 | 0,361371825 | 5,497802967 | 3,85E-08 | 1,16E-05 | Ensembl Transcript | 5579990 | Gm29284 |
| ENSMUST00000135184 | 917,9578437 | 2,014524908 | 0,410387871 | 4,9088315 | 9,16E-07 | 0,00015037 | Ensembl Transcript | 97243 | Mx1 |
| ENSMUST00000219513 | 3,498371804 | 2,036053441 | 0,635476624 | 3,203978503 | 0,001355426 | 0,029378873 | Ensembl Transcript | 1924652 | 9530018F02Rik |
| ENSMUST00000216328 | 12,42620999 | 2,079050901 | 0,358858911 | 5,793505019 | 6,89E-09 | 2,61E-06 | Ensembl Transcript | 2177482 | Olfr99 |
| ENSMUST00000169041 | 4,189082127 | 2,08118653 | 0,680317712 | 3,059139123 | 0,00221974 | 0,041242498 | Ensembl Transcript | 1926156 | Misp |
| ENSMUST00000166912 | 7,618235306 | 2,128187438 | 0,568756123 | 3,741827736 | 0,000182687 | 0,00693693 | Ensembl Transcript | 3648476 | Phf11c |
| ENSMUST00000050011 | 9,006660736 | 2,142721784 | 0,676498275 | 3,167372131 | 0,001538233 | 0,032046891 | Ensembl Transcript | 5663439 | Gm43302 |
| ENSMUST00000172538 | 103,507649 | 2,152054722 | 0,187414398 | 11,48286764 | 1,61E-30 | 4,75E-27 | Ensembl Transcript | 2442805 | H2-T-ps |

| | | | | | | | | | |
|---|---|---|---|---|---|---|---|---|---|
| ENSMUST00000020969 | 1133,809143 | 2,187028407 | 0,398908996 | 5,482524661 | 4,19E-08 | 1,21E-05 | Ensembl Transcript | 99830 | Cmpk2 |
| ENSMUST00000076249 | 130,7510067 | 2,190631018 | 0,502105631 | 4,362888771 | 1,28E-05 | 0,001184968 | Ensembl Transcript | 3698419 | Ifit3b |
| ENSMUST00000007156 | 6,720168556 | 2,347583323 | 0,673614945 | 3,485052315 | 0,000492041 | 0,014387157 | Ensembl Transcript | 892023 | Klk1b11 |
| ENSMUST00000178990 | 3,936634998 | 2,35607628 | 0,789580938 | 2,983957902 | 0,002845459 | 0,049158219 | Ensembl Transcript | 3704307 | Proscos |
| ENSMUST00000149829 | 3088,092983 | 2,361508157 | 0,494684324 | 4,773767924 | 1,81E-06 | 0,000269773 | Ensembl Transcript | 99449 | Ifit2 |
| ENSMUST00000143783 | 3,710122663 | 2,36693359 | 0,681544756 | 3,472895317 | 0,000514876 | 0,01471032 | Ensembl Transcript | 96785 | Lhx2 |
| ENSMUST00000020970 | 3785,996356 | 2,383575095 | 0,41359703 | 5,763037255 | 8,26E-09 | 3,04E-06 | Ensembl Transcript | 1929628 | Rsad2 |
| ENSMUST00000113760 | 1321,489174 | 2,467360085 | 0,263713835 | 9,356202674 | 8,27E-21 | 1,11E-17 | Ensembl Transcript | 95958 | H2-T24 |
| ENSMUST00000173128 | 15,46820892 | 2,470314625 | 0,34873697 | 7,083604086 | 1,40E-12 | 1,04E-09 | Ensembl Transcript | 5011869 | Gm19684 |
| ENSMUST00000024757 | 55,27353245 | 2,736451447 | 0,323999362 | 8,445854425 | 3,02E-17 | 2,48E-14 | Ensembl Transcript | 2682634 | Enpp4 |
| ENSMUST00000173900 | 27,63631049 | 2,901044889 | 0,289538989 | 10,01953104 | 1,25E-23 | 3,08E-20 | Ensembl Transcript | 1925060 | A930015D03Rik |
| ENSMUST00000024805 | 1,594194877 | 3,351283147 | 1,00029478 | 3,350295548 | 0,000807254 | 0,0201143 | Ensembl Transcript | 2385908 | Cpne5 |
| ENSMUST00000098827 | 2,478841449 | 3,423041529 | 1,05660066 | 3,239673851 | 0,001196665 | 0,027102643 | Ensembl Transcript | 3641851 | Gm10684 |
| ENSMUST00000018610 | 4,130986481 | 3,502738894 | 0,786202663 | 4,455262059 | 8,38E-06 | 0,000859496 | Ensembl Transcript | 97361 | Nos2 |
| ENSMUST00000122936 | 1,539990546 | 3,80004462 | 1,239433929 | 3,065951747 | 0,002169783 | 0,040672422 | Ensembl Transcript | 1918253 | Prss41 |
| ENSMUST00000024928 | 1,451322178 | 3,909398516 | 1,282950926 | 3,04719256 | 0,002309897 | 0,042384455 | Ensembl Transcript | 1916698 | Prss21 |
| ENSMUST00000094972 | 1,691135048 | 4,372539239 | 1,195450155 | 3,65765082 | 0,000254537 | 0,008623333 | Ensembl Transcript | 107668 | Ifna1 |
| ENSMUST00000172895 | 4,417869081 | 5,572262755 | 0,914832404 | 6,091020312 | 1,12E-09 | 4,60E-07 | Ensembl Transcript | 3779814 | Gm8810 |
| ENSMUST00000021822 | 16,66415962 | 22,3359845 | 4,857694332 | 4,598062986 | 4,26E-06 | 0,000524909 | Ensembl Transcript | 109278 | Ogn |

**Table EV2.** Identity of down-regulated phosphosites (significantly enriched) that are specific to S/T-P motif in p38γ/δ$^{-/-}$ and p38γ/δKIKO macrophages compared to WT cells.

| **p38γ/δ-/- v WT comparison** | | | |
|---|---|---|---|
| Protein | Gene | Position | Sequence |
| Nuclear factor NF-kappa-B p100/p52 subunit | Nfkb2 | 425 | SRRDTDAGEGAEEPR**TP**PEAPQGEPQALDTL |
| Myocyte-specific enhancer factor 2D | Mef2d | 437 | TVTTHPHISIKSEPV**SP**SRERSPAPPPPAVF |
| Probable C-mannosyltransferase DPY19L1 | Dpy19l1 | 46 | SEVDAGELGSERTPP**SP**GRRGAAGRKGPRAG |
| Ataxin-2-like protein | Atxn2l | 407 | PGSEARGINGGPSRM**SP**KAQRPLRGAKTLSS |

| **p38γ/δKIKO v WT comparison** | | | |
|---|---|---|---|
| Protein | Gene | Position | Sequence |
| Putative adenosylhomocysteinase 3 | Ahcyl2 | 109 | HHQHQRHRDGGEALV**SP**DGTVTEAPRTVKKQ |
| Myocyte-specific enhancer factor 2D | Mef2d | 437 | TVTTHPHISIKSEPV**SP**SRERSPAPPPPAVF |
| SLAM family member 7 | Slamf7 | 313 | PNTFYSTVQIPKVVK**SP**SSLPAKPLVPRSLS |
| RUN and FYVE domain-containing protein 1 | Rufy1 | 52 | ESGEAFEIVDRSQLP**SP**GELRSASRPRVADS |
| Stathmin | Stmn1 | 25 | ELEKRASGQAFELIL**SP**RSKESVPDFPLSPP |
| Tumor protein D52 | Tpd52 | 170 | SVGSVITKKLEDVKN**SP**TFKSFEEKVENLKS |
| Uncharacterized protein KIAA1522 | Kiaa1522 | 909 | IFSKGTKKLQLERPV**SP**EAQADLQRNLVAEL |
| Histone-lysine N-methyltransferase 2D | Kmt2d | 2299 | FKAPLTPRASQVEPQ**SP**GLGLRAQEPPPAQA |
| ADP-ribosylation factor-like protein 6-interacting protein | Arl6ip6 | 80 | SETKRAPLLPRVGDG**SP**VLPDKRNGIFPATA |
| Lamin-B1 | Lmnb1 | 21 | PVQQQRAGSRASAPA**TP**LSPTRLSRLQEKEE |
| MAP kinase-interacting serine/threonine-protein kinase | Mknk1 | 344 | QHPWVQGQAPERGLP**TP**QVLQRNSSTMDLTL |

**Table EV3. Sites on MEF2D phosphorylated by p38α or p38δ *in vitro*.** Human GST-MEF2D was incubated in a phosphorylation reaction mix containing Mg-ATP in the absence (control) or the presence of p38α or p38δ, and subjected to SDS-PAGE. Phosphorylated MEF2D was excised from the gel, digested with trypsin and peptides separated as indicated in Materials and Method. Phosphorylated residues were identified by LC-MS/MS. Peptides containing the phosphorylated residues are shown. Phosphorylated residues are indicated as pS in red, deaminated residues are indicated as (deam) in blue.

| Protein phospho-site (Q14814) | Phosphopeptide | control | p38α | p38δ |
|---|---|---|---|---|
| S98 | KGFN(deam)GCDpSPEPDGEDSLEQSPLLEDK | X | X | |
| | KGFNGCDpSPEPDGEDSLEQSPLLEDK | | | |
| | GFNGCDpSPEPDGEDSLEQSPLLEDK | | | |
| | GFNGCDpSPEPDGEDSLEQSPLLEDKYR | | | |
| S121 | RApSEELDGLFR | X | | X |
| S180 | LLpSPQQPALQR | X | X | X |
| S192 | NSVpSPGLPQR | | X | |
| S231 | ApSPGLLPVANGNSLNK | X | X | X |
| | ApSPGLLPVAN(deam)GNSLNK | | | |
| | ApSPGLLPVAN(deam)GN(deam)SLNK | | | |
| S251 | VIPAKpSPPPPTHSTQLGAPSR | | X | X |
| | pSPPPPTHSTQLGAPSR | | | |
| S275 | VITpSQAGK | | | X |
| S444 | SEPVpSPSRER | | | X |
| S450 | ERpSPAPPPPAVFPAAR | | X | X |
| S472 | PEPGDGLSpSPAGGSYETGDR | | | X |

**Table EV4.** Information about the antibodies used in this study.

| Antibody | Name | Provider | Method | Dilution |
|---|---|---|---|---|
| P-ERK1/2 | Phospho-p44/42 MAPK (Erk1/2) (Thr202/Tyr204) Antibody | Cell Signaling | WB | 1/2000 |
| ERK1/2 | p44/42 MAPK (Erk1/2) Antibody | Cell Signaling | WB | 1/1000 |
| TPL2 | Cot (M-20) | Santa Cruz Biotechnology, Inc. | WB | 1/1000 |
| ABIN2 | ABIN2 antibody | S.C. Ley laboratory | WB | 2 μg/ml |
| P-p38 | Phospho-p38 MAP Kinase (Thr180/Tyr182) Antibody | Cell Signaling | WB | 1/1000 |
| p38α | p38α (C-20) | Santa Cruz Biotechnology, Inc. | WB | 1/1000 |
| p38γ | p38γ (GST-SAPK3) S524A 1st Bleed | DSTT* | WB | 1/1000 |
| p38δ | p38δ (GST-SAPK4) S526A 3rd Bleed | DSTT* | WB | 1/1000 |
| P-JNK1/2 | Rabbit (polyclonal) Anti-JNK1&2 [pTpY183/185] Phosphospecific Antibody, Unconjugated | Biosource | WB | 1/1000 |
| JNK1/2 | SAPK/JNK | Cell Signaling | WB | 1/1000 |
| P-p105 | Phospho-NF-κB p105 (Ser933) (18E6) Rabbit mAb | Cell Signaling | WB | 1/1000 |
| p105 | NF-κB1 p105 Antibody | Cell Signaling | WB | 1/1000 |
| hDlg | hDlg (SAP97) antibody | DSTT* | WB | 1/1000 |
| P-hDlg (S158) | Phospho-hDlg (SAP97) $Ser^{158}$ | DSTT* | WB | 1/1000 |
| hDlg | hDlg (SAP97) antibody | DSTT* | IP | 1 μg/IP |
| p38γ | p38γ (SAPK3 C-Terminal) [KPPRNLGARVPKETA] | DSTT* | IP | 2 μg/IP |
| CD45 | CD45 Monoclonal Antibody (30-F11), APC #17-0451-82 | eBioscience™ | FAC | 1/100 |
| F4/80 | F4/80 Monoclonal Antibody (BM8) FITC #11-4801-85 | eBioscience™ | FAC | 1/100 |
| Ly6G | PE Rat Anti-Mouse Ly-6G Clone 1A8 (RUO) #551461 | BD Bioscience | FAC | 1/50 |

* DSTT (Dundee): Division of Signal Transduction Therapy; University of Dundee (Dundee, UK)

**Table EV5:** Primer sequences used for gene expression

| Gene | Forward (5´-3´) | Reverse (5´-3´) |
|---|---|---|
| TNFα | TTGAGATCCATGCCGTTG | CTGTAGCCCACGTCGTAGC |
| IL-1β | TGGTGTGTGACGTTCCCATT | CAGCACGAGGCTTTTTTGTTG |
| IL-6 | GAGGATACCACTCCCAACAGACC | AAGTGCATCATCGTTGTTCATACA |
| IFNβ | TCAGAATGAGTGGTGGTTGC | GACCTTTCAAATGCAGTAGATTCA |
| iNOS | CAGCTGGGCTGTACAAACCTT | CATTGGAAGTGAAGCGTTTCG |
| IL12 p35 | CCACCCTTGCCCTCCTAAA | GGCAGCTCCCTCTTGTTGTG |
| CCL5 | ATATGGCTCGGACACCACTC | TTCTTCGAGTGACAAACACG |
| CXCL9 | TGCACGATGCTCCTGCA | AGGTCTTTGAGGGATTTGTAGTGG |
| CXCL10 | GGATCCCTCTCGCAAGGA | ATCGTGGCAATGATCTCAACA |
| KC (CXCL1) | CCTTGACCCTGAAGCTCCCT | CGGTGCCATCAGAGCAGTCT |
| MCP1 (CCL2) | TCTGGGCCTGCTGTTCACA | TTGGGATCATCTTGCTGGTG |
| MIP2 (CXCL2) | CCTGGTTCAGAAAATCATCCA | CTTCCGTTGAGGGACAGC |
| c-Jun | GCACATCACCACTACACCGA | GGGAAGCGTGTTCTGGCTTAT |
| CD14 | CATTTGCATCCTCCTGGTTTCTGA | GAGTGGTTTTCCCCTTCCGTGTG |
| HDAC7 | CGCAGCCAGTGTGAGTGTCT | AGTGGGTTCGTGCCGTAGAG |
| β-actin | AAGGAGATTACTTGCTCTGGCTCCTA | ACTCATCGTACTCCTGCTTGCTGAT |
